# Proton co-transport and voltage dependence enforce unidirectional metal transport in an Nramp transporter

**DOI:** 10.1101/402412

**Authors:** Aaron T. Bozzi, Lukas B. Bane, Christina M. Zimanyi, Rachelle Gaudet

## Abstract

Secondary transporters harness electrochemical energy to move substrate through structurally-enforced co-substrate “coupling”. We untangle the “proton-metal coupling” behavior by a Natural resistance-associated macrophage protein (Nramp) transporter into two distinct phenomena: ΔpH stimulation of metal transport and metal stimulation of proton co-transport. Surprisingly, metal type dictates co-transport stoichiometry, leading to manganese-proton symport but cadmium uniport. Additionally, the membrane potential affects both the kinetics and thermodynamics of metal transport. A conserved salt-bridge network near the metal-binding site imparts voltage dependence and enables proton co-transport, properties that allow this Nramp transporter to maximize metal uptake and prevent deleterious back-transport of acquired metals. We provide a new mechanistic model for Nramp metal-proton symport in which, in addition to substrate gradients determining directionality as in canonical secondary transport, synergy between protein structure and physiological voltage enforces unidirectional substrate movement. Our results illustrate a functional advantage that arises from deviations from the traditional model of symport.

---

Cells commit significant ATP expenditure to maintain ionic gradients across membranes, a form of energy storage. Selective membrane permeability also generates a stable charge imbalance that produces a membrane voltage, generally negative. Secondary transporters harness these electrochemical gradients by coupling the energetically-favorable movement of abundant ions (Na^+^, K^+^, H^+^, Cl^-^) to the (often uphill) movement of substrates (Forrest et al., 2011; Gadsby, 2009; LeVine et al., 2016; Shilton, 2015). Tight coupling mechanisms prevent uniport events: “futile cycles” that dissipate the driving ion gradients and, even more deleterious, the back-transport of the primary substrate down its concentration gradient (LeVine et al., 2016).

The Nramp (Natural resistance-associated macrophage protein) family of transporters import divalent transition metal ions, essential micronutrients serving as cofactors to myriad metabolic enzymes. Prokaryotic Nramps perform high-affinity Mn^2+^ scavenging (Cellier, 2012; Ma et al., 2009), while in eukaryotes, including humans, Nramps are essential to iron dietary uptake, iron trafficking to and recycling from erythrocytes, and the metal-withdrawal innate immune defense (Abbaspour et al., 2014; Andrews, 2008; Coffey and Ganz, 2017).

Nramps are thought to couple proton transport to metal substrate import (Chen et al., 1999a; Gunshin et al., 1997)—primarily Mn^2+^ and Fe^2+^, but also the biologically useful metals Co^2+^, Ni^2+^, and Zn^2+^ (Ehrnstorfer et al., 2014; Illing et al., 2012; Picard et al., 2000); and toxic heavy metals Cd^2+^, Pb^2+^, and Hg^2+^ (Bressler et al., 2004; Vazquez et al., 2015). Given typically low environmental metal ion concentrations, this coupling may make the net transport thermodynamically favorable. However, the substantial negative membrane potential (which favors cation entry) and the plethora of intracellular metal-binding proteins and small molecules (which likely act as a sink) may make metal entry alone energetically-downhill in many cases. Early electrophysiological studies of Nramps showed variable metal-proton transport stoichiometries depending on pH and membrane potential, with significant proton uniport, suggesting loose coupling (Chen et al., 1999a; Mackenzie et al., 2006; Nelson et al., 2002). Furthermore, multiple Nramp homologs exhibit voltage-and pH-dependence of transport (Chen et al., 1999a; Gunshin et al., 1997; Mackenzie et al., 2007; Xu et al., 2004). Nramps have a common secondary-transporter fold (Ehrnstorfer et al., 2014) and an alternating-access conformational cycle (Bozzi et al., 2016b; Bozzi et al., 2018; Ehrnstorfer et al., 2017). Several conserved residues are important for substrate selectivity (Bozzi et al., 2016a; Bozzi et al., 2018) and proton-metal coupling (Ehrnstorfer et al., 2017; Pujol-Gimenez et al., 2017).

We use the *Deinococcus radiodurans* (Dra)Nramp homolog, for which we determined snapshots of the structure in multiple conformations (Bozzi et al., 2016b; Bozzi et al., 2018), to demonstrate how it harnesses the membrane potential to both accelerate metal import and prevent deleterious metal export. We further demonstrate how the Nramp “proton-metal coupling” phenotype encompasses two distinct phenomena: ΔpH stimulates metal transport, and metal stimulates proton co-transport, with some metal substrates surprisingly undergoing uniport—metal transport without accompanying proton(s)—rather than the expected symport. We show how a conserved salt-bridge network allosterically imparts voltage dependence and forms the pathway for proton co-transport. Our results mechanistically explain how an Nramp provides unidirectional transport for its transition metal substrates through its many deviations from the canonical model for symport.

## Results

### The membrane potential drives DraNramp metal transport

Our investigation of the role of transmembrane voltage in DraNramp’s mechanism stems from trying to reconcile an apparent inconsistency between *in vivo* and *in vitro* protein behavior. Purified DraNramp reconstituted into proteoliposomes (Figure 1A and Figure 1—figure supplement 1A) with large metal substrate gradients transported Cd^2+^—a toxic metal known to be a good Nramp substrate (Courville et al., 2008; Illing et al., 2012)—but not the biological substrates Mn^2+^ or Fe^2+^, or similar metals like Zn^2+^ or Co^2+^ (Figure 1B and Figure 1—figure supplement 1D). In contrast DraNramp robustly transported Co^2+^ when expressed in *Escherichia coli* (Figure 1—figure supplement 1B).

**Figure 1.**
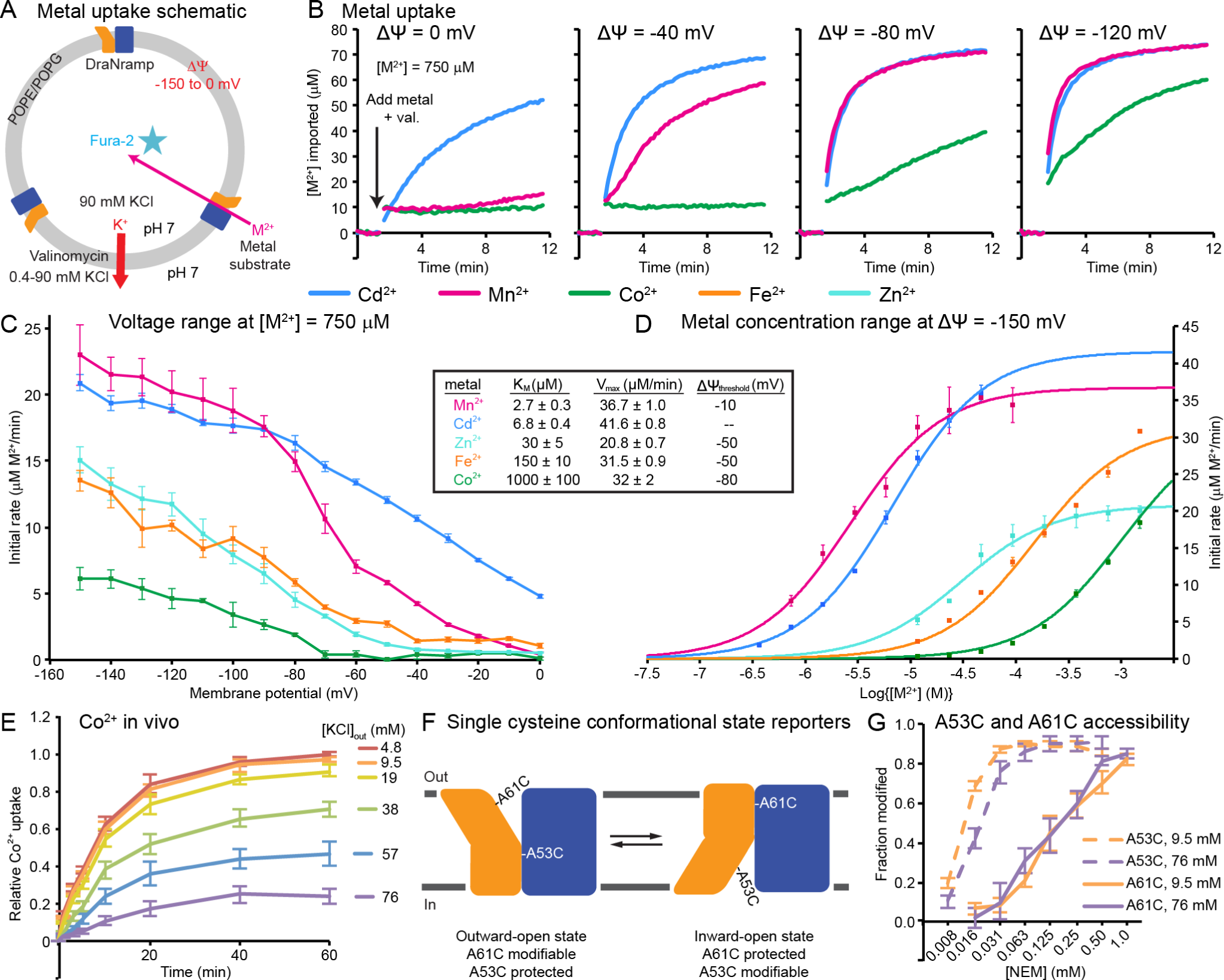
Metal transport shows strong voltage dependence. (A) Schematic of proteoliposome assay for metal transport at variable membrane potentials (ΔΨ) established by using K^+^ gradients and valinomycin. (B) Representative time traces (n = 8) of Cd^2+^, Mn^2+^, and Co^2+^ transport at different ΔΨs; only Cd^2+^ is transported at ΔΨ = 0 mV. (C) Average initial transport rate ± S.E.M. (n = 4) versus membrane potential. Each metal except Cd^2+^ has a characteristic threshold negative voltage for transport to occur: −10 mV for Mn^2+^, −50 mV for Fe^2+^ and Zn^2+^, and −80 mV for Co^2+^. (D) Average initial transport rate ± S.E.M. (n = 2-3) versus metal concentration. Comparison of panels C and D (table inset) show that higher-magnitude threshold voltages correlate with lower apparent affinity (K_M_), indicating that voltage dependence may be enforced through the metal-binding site. Errors in K_M_ and V_max_ encompass the uncertainty of fit (shown as curved lines) to data. (E) Time course of Co^2+^ uptake shows that high extracellular [KCl] depresses Co^2+^ uptake in *E. coli*. (F) Complementary single-cysteine reporters to assess conformational cycling of DraNramp. (G) Fraction of modified cysteine versus NEM concentration. Inward (A53C) and outward (A61C) reporters both label even at high [KCl]_out_ that depresses Co^2+^ uptake. Conformational-locking thus cannot explain reduced Co^2+^ transport by DraNramp under these conditions. Data in E and G are averages ± S.E.M. (n = 4). See also Figure 1—figure supplement 1.

Bacteria, including *E. coli*, maintain a membrane potential (ΔΨ) between −140 and −220 mV (Bot and Prodan, 2010), which greatly influences ion transport thermodynamics. When we applied voltage across proteoliposome membranes (see Methods), DraNramp transported Mn^2+^ at −40 mV or below (Figure 1B), and Fe^2+^, Zn^2+^, and Co^2+^ at −80 mV (Figure 1B and Figure 1—figure supplement 1D), with notable acceleration at −120 mV (Figure 1B and Figure 1—figure supplement 1D). Indeed, by measuring at 10 mV increments, we saw metal-specific threshold voltages for metal transport with Mn^2+^, followed by Zn^2+^ and Fe^2+^, then Co^2+^ requiring progressively larger potential differences to observe metal transport, while Cd^2+^ was transported across the whole measured range (Figure 1C and Figure 1—figure supplement 1E). These trends are similar to the voltage-dependence of mammalian Nramp2 Fe^2+^ transport observed via electrophysiology (Gunshin et al., 1997; Mackenzie et al., 2006; Pujol-Gimenez et al., 2017; Sacher et al., 2001).

We next measured transport across a range of metal concentrations at a physiologically-relevant ΔΨ = −150 mV (Figure 1D) to determine Michaelis-Menten constants. The apparent substrate affinity (KM) at −150 mV correlated with the requisite threshold voltage (Figure 1C): Mn^2+^ and Cd^2+^ had highest affinity and the lowest magnitude threshold ΔΨ, Co^2+^ had the lowest affinity and highest magnitude threshold ΔΨ, and Zn^2+^ and Fe^2+^ were intermediate for both. Our KM values are consistent with *in vivo* Co^2+^ uptake data (Figure 1—figure supplement 1F) and of similar magnitude to previously reported values for *in vivo* Cd^2+^ and Mn^2+^ transport by the *E. coli* Nramp homolog (Courville et al., 2008; Haemig and Brooker, 2004b). The much higher apparent affinity (KM < 10 μM) for Mn^2+^ and Cd^2+^ than other divalent metal substrates is consistent with trends first observed in *E. coli* Nramp (Makui et al., 2000), but contrasts with human Nramp2 (DMT1), which has affinities for Mn^2+^, Fe^2+^, Co^2+^, Zn^2+^, and Cd^2+^ all within an order of magnitude in the low μM range (Illing et al., 2012). These results suggest that these bacterial homologs have higher specificity for their intended substrate, Mn^2+^, with incidental high Cd^2+^ affinity. Next, to recapitulate the voltage-dependence trend *in vivo*, we varied external [K^+^], which likely perturbed the membrane potential of *E. coli*, as they maintain a significant cytoplasmic [K^+^] (Shabala et al., 2009). High [K^+^] indeed drastically decreased DraNramp Co^2+^ transport (Figure 1E and Figure 1—figure supplement 1G; see Figure 5—figure supplement 2 below for comparison of DraNramp point mutants).

As one explanation for our results, the membrane potential might perturb DraNramp’s gross conformational landscape, with the outward-facing state relatively destabilized at lower magnitude voltages, as seen with the structurally-related SGLT1 transporter (Loo et al., 1998). We therefore measured the accessibility to N-ethylmaleimide (NEM) modification of single-cysteine reporters that are only exposed to bulk solvent in either the inward-open (A53C) or outward-open (A61C) states (Bozzi et al., 2016b; Bozzi et al., 2018) (Figure 1F). However, although high extracellular [K^+^] greatly impaired Co^2+^ transport, it had little effect on the sampling of either the inward-facing or outward-facing states (Figure 1G and Figure 1—figure supplement 1H). We conclude that DraNramp voltage dependence arises from more nuanced mechanistic changes. This is more consistent with the observed substrate-specific threshold voltages, as a shift in conformational equilibrium that completely inhibits transport of all metal does not account for this substrate specificity.

Our consistent *in vivo* and *in vitro* findings indicate the reconstituted transporter does indeed function the same as in the native membrane, and that transmembrane voltage—the missing piece in our initial reconstitution attempts—is a critical variable controlling metal transport kinetics.

### Proton co-transport is substrate-specific in DraNramp

Despite their similar KM and Vmax at physiological ΔΨ, Mn2+ and Cd2+ behave very differently at voltages near zero. Indeed, when extending our liposome assay to values of ΔΨ > 0, which thermodynamically disfavors cation entry, DraNramp still transported Cd^2+^—but not Mn^2+^— down its concentration gradient even at +50 mV (Figure 2—figure supplement 1A-C), hinting at a mechanistic difference between the two metals.

In addition to ΔΨ’s importance, the well-documented requirement of many Nramp homologs for low pH to stimulate metal transport (Chen et al., 1999a; Gunshin et al., 1997; Makui et al., 2000) suggests that ΔpH is also a critical variable for DraNramp. We therefore compared the pH dependence of transport for both Cd^2+^ and Mn^2+^ (Figure 2A). While lower external pH moderately enhanced Cd^2+^ transport (Figure 2B), it greatly accelerated Mn^2+^ transport—even enabling transport at 0 mV (Figure 2C). Therefore Mn^2+^ transport—but not Cd^2+^—may require thermodynamically favorable proton entry, either through a negative membrane potential or a pH gradient.

**Figure 2.**
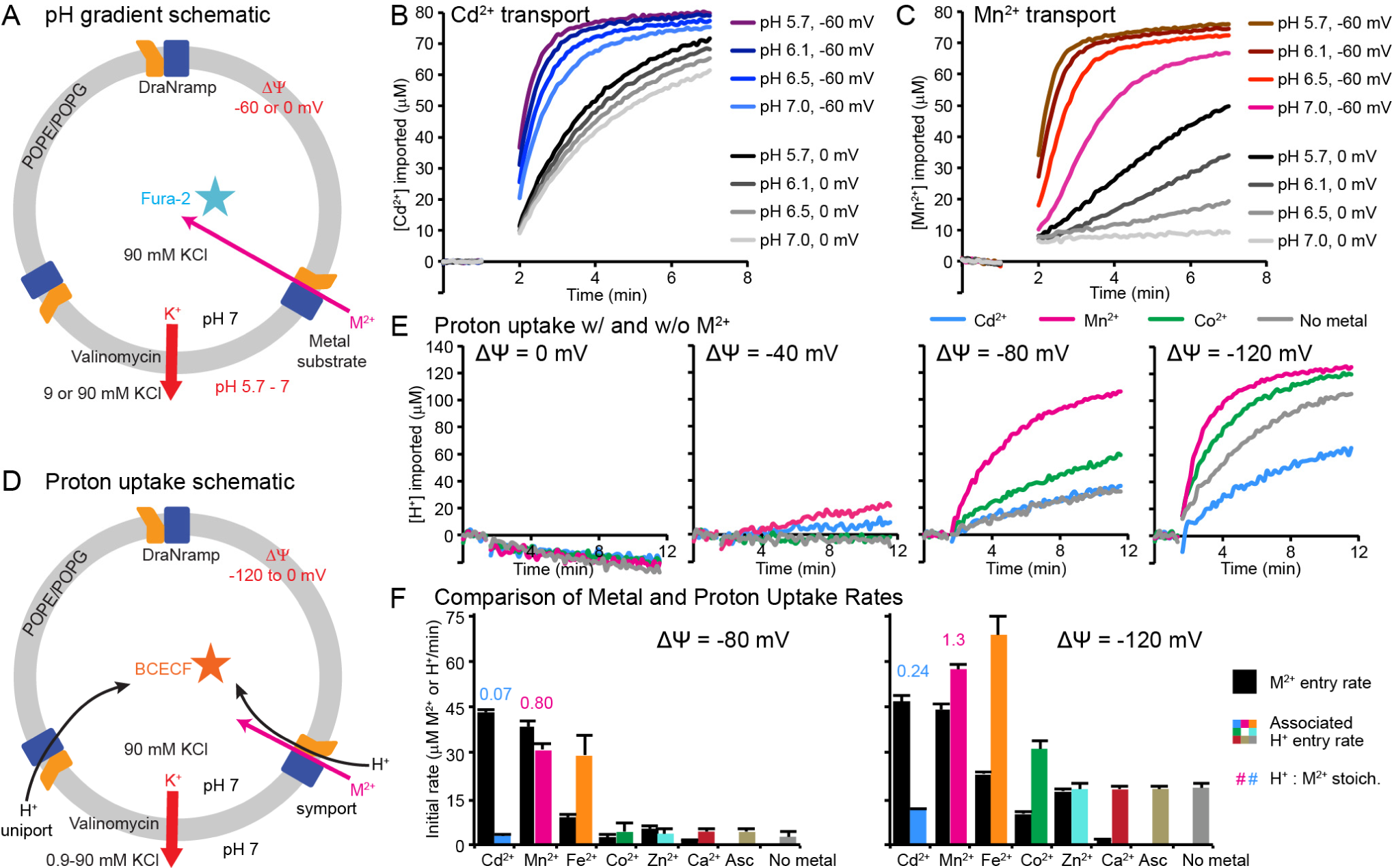
pH stimulation is distinct from proton co-transport. (A) Schematic for assessing ΔpH effect on metal transport in proteoliposomes. (B-C) Representative time traces (n ≥ 4) of Cd^2+^ (B) or Mn^2+^ (C) uptake into proteoliposomes. ΔpH accelerated Cd^2+^ and Mn^2+^ transport (at ΔΨ = −60 mV), and enabled Mn^2+^ uptake when ΔΨ = 0. (D) Schematic for detecting H^+^ influx into proteoliposomes. (E) Representative time traces (n = 8) of H^+^ uptake into proteoliposomes. Negative ΔΨ drove DraNramp H^+^ import. Mn^2+^ and Co^2+^ stimulated H^+^ entry, while Cd^2+^ did not, and instead reduced H^+^ influx at ΔΨ = −120 mV. Except for the reporter dye, conditions were identical to those in Figure 1B. (F) Initial metal (black) and H^+^ (color) uptake rates show that Mn^2+^, Fe^2+^, and Co^2+^ transport stimulated H^+^ influx above its basal no-metal rate, while Cd^2+^ and Zn^2+^ transport did not. Stoichiometry ratios (numbers above bars) were calculated for Cd^2+^ and Mn^2+^ and were approximately 1:1 for Mn^2+^/H^+^ symport. Data are averages ± S.E.M. (n ≥ 4). See also Figure 2—figure supplement 1.

To test this hypothesis, we measured H^+^ transport by DraNramp (Figure 2D) at a range of voltages in the absence or presence of metal substrate (Figure 2E and Figure 2—figure supplement 1D). While DraNramp efficiently transported all tested metals (except Ca^2+^) at −120 mV and to some degree at −80 mV, only Mn^2+^, Fe^2+^, and Co^2+^ stimulated H^+^ influx above basal rates (Figure 2F). Despite robust Cd^2+^ uptake across all potentials, Cd^2+^ failed to stimulate significant H^+^ transport. Indeed at −120 mV, where H^+^ uniport—uptake in the absence of added metal—became significant, Cd^2+^ noticeably reduced the H^+^ influx rate (Figure 2E-F). DraNramp’s two best substrates, which should nearly saturate the metal-binding site at −80 and −120 mV, gave initial rate stoichiometries of approximately 1 H^+^: 1 Mn^2+^ and 0 H^+^: 1 Cd^2+^ (Figure 2F), confirming that metal-proton co-transport is indeed substrate-specific.

### Voltage and pH both shape DraNramp’s free energy landscape

To better understand the effects of transmembrane voltage and pH gradients on DraNramp transport, we determined Michaelis-Menten parameters for Mn^2+^ and Cd^2+^ transport kinetics in the absence or presence of a pH gradient at three ΔΨ values (Figure 3A-B) and used our results to sketch reaction coordinate diagrams for Cd^2+^ and Mn^2+^ transport (Figure 3C-D). For both metals at both pHs, a more negative ΔΨ increased transport efficiency by improving both KM and Vmax. Strikingly, in the absence of ΔpH, hyperpolarizing ΔΨ from −50 to −150 mV decreased the KMs for Cd^2+^ and Mn^2+^ more than 20- and 300-fold respectively. Thus, in addition to its impact on ion uptake thermodynamics, ΔΨ also shapes the free energy landscape that dictates DraNramp transport kinetics (Figure 3C-D) and thus likely influences metal binding and other steps in the transport mechanism.

**Figure 3.**
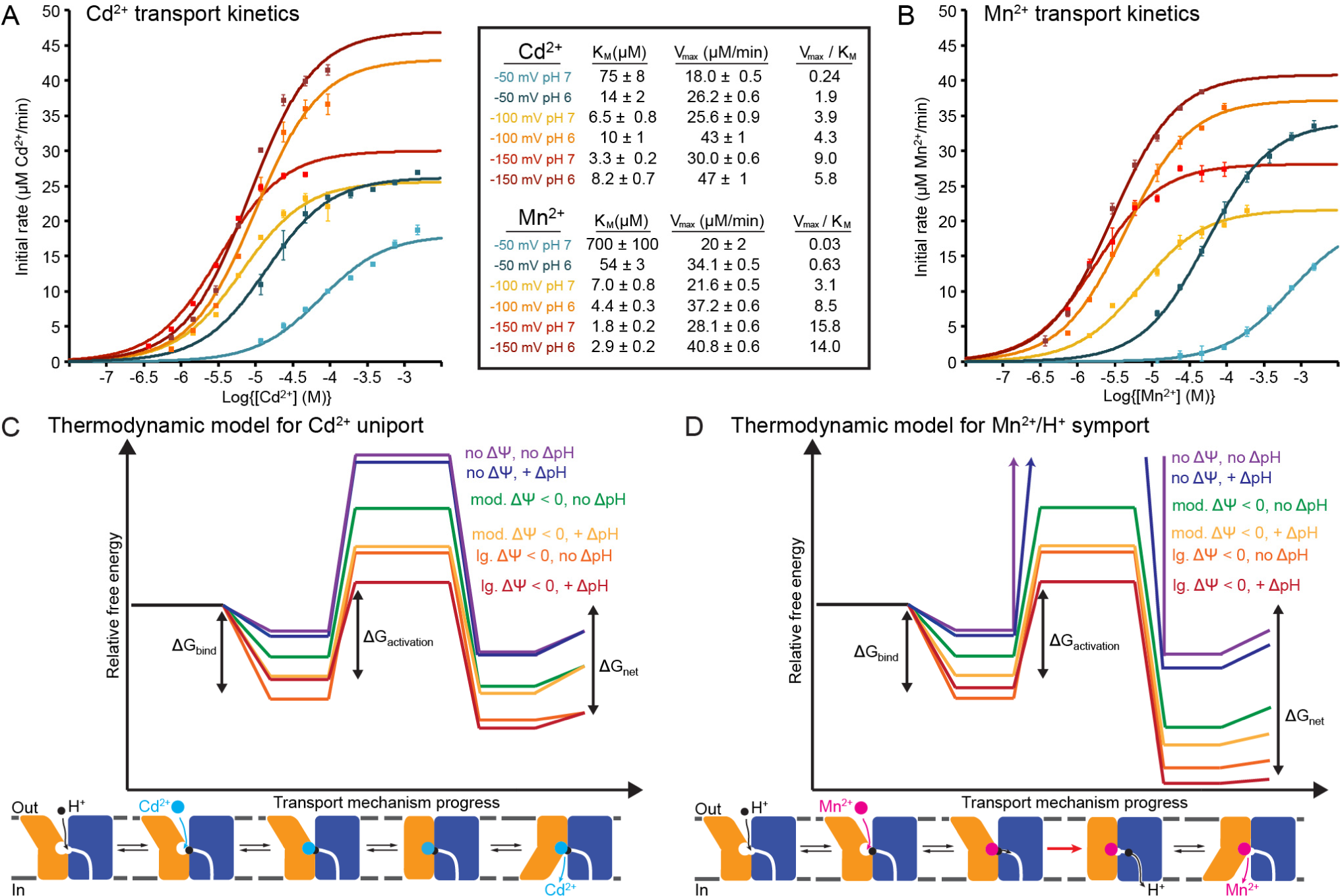
pH and voltage govern DraNramp transport kinetics. (A-B) Dose-response curves for Cd^2+^ (A) and Mn^2+^ (B) transport at different external pH and ΔΨ values. Data are averages ± S.E.M. (n = 3). Errors in K_M_ and V_max_ reported in the inset table encompass the uncertainty of the fit (shown as curved lines) to the data. (C-D) Proposed free energy landscapes of Cd^2+^ (C) or Mn^2+^ (D) transport based on transport data in (A-B) and Figures 1-2.

While in all six cases, ΔpH increased Vmax, the effects on KM were mixed (Figure 3A-B). Indeed, ΔpH enhanced apparent affinity for both metals at lower magnitude ΔΨ but reduced apparent affinity at higher magnitude ΔΨ. Generally, ΔΨ and ΔpH are more synergistic for Mn^2+^ and more antagonistic for Cd^2+^, consistent with the H^+^ co-transport requirement for Mn^2+^ but not Cd^2+^ (Figure 2F). In addition, our data indicate that an insurmountable kinetic barrier precludes even downhill Mn^2+^—but not Cd^2+^—transport in the absence of a ΔΨ and/or ΔpH favorable to H^+^ transport (Figure 3C-D). Taken together with the fact that ΔpH affected the transport kinetics of both metals, this indicates that “proton-metal coupling” in Nramps comprises at least two distinct phenomena: (i) low pH/ΔpH stimulates metal transport, and (ii) H^+^ co-transport is required for certain metal substrates. Below, using insights gleaned from DraNramp’s structures and mutagenesis experiments we begin to mechanistically explain these surprising results.

### DraNramp metal-binding site abuts an extended salt-bridge network

Our structures of DraNramp in multiple conformational states (Bozzi et al., 2016b; Bozzi et al., 2018) provide a framework to understand the mechanistic details of voltage dependence, ΔpH stimulation, and proton co-transport. DraNramp’s 11 α-helical transmembrane (TM) segments adopt the common LeuT-fold (Boudker and Verdon, 2010; Forrest et al., 2011; Shi, 2013; Yamashita et al., 2005) (Figure 4A). Conserved residues D56, N59, and M230 coordinate metal substrate throughout the transport process (Bozzi et al., 2018; Ehrnstorfer et al., 2014) (Figure 4B). Below the binding site, conserved H237 on TM6b lines the metal’s exit route to the cytoplasm (Bozzi et al., 2018) (Figure 4C and Figure 4—figure supplement 1A). Adjacent to D56, a conserved network of charged and protonatable residues lead into the structurally-rigid cluster formed by TMs 3, 4, 8, and 9 to provide a parallel ∼20 Å H^+^-transport pathway towards the cytoplasm (Bozzi et al., 2018) (Figure 4C-D and Figure 4—figure supplement 1A-B). These residues form a sequence of three interacting pairs: H232-E134, D131-R353, and R352-E124. This salt-bridge network is unique to the Nramp clade in the LeuT family of structurally- and evolutionarily-related transporters (Bozzi et al., 2018; Cellier, 2012; Cellier, 2016).

**Figure 4.**
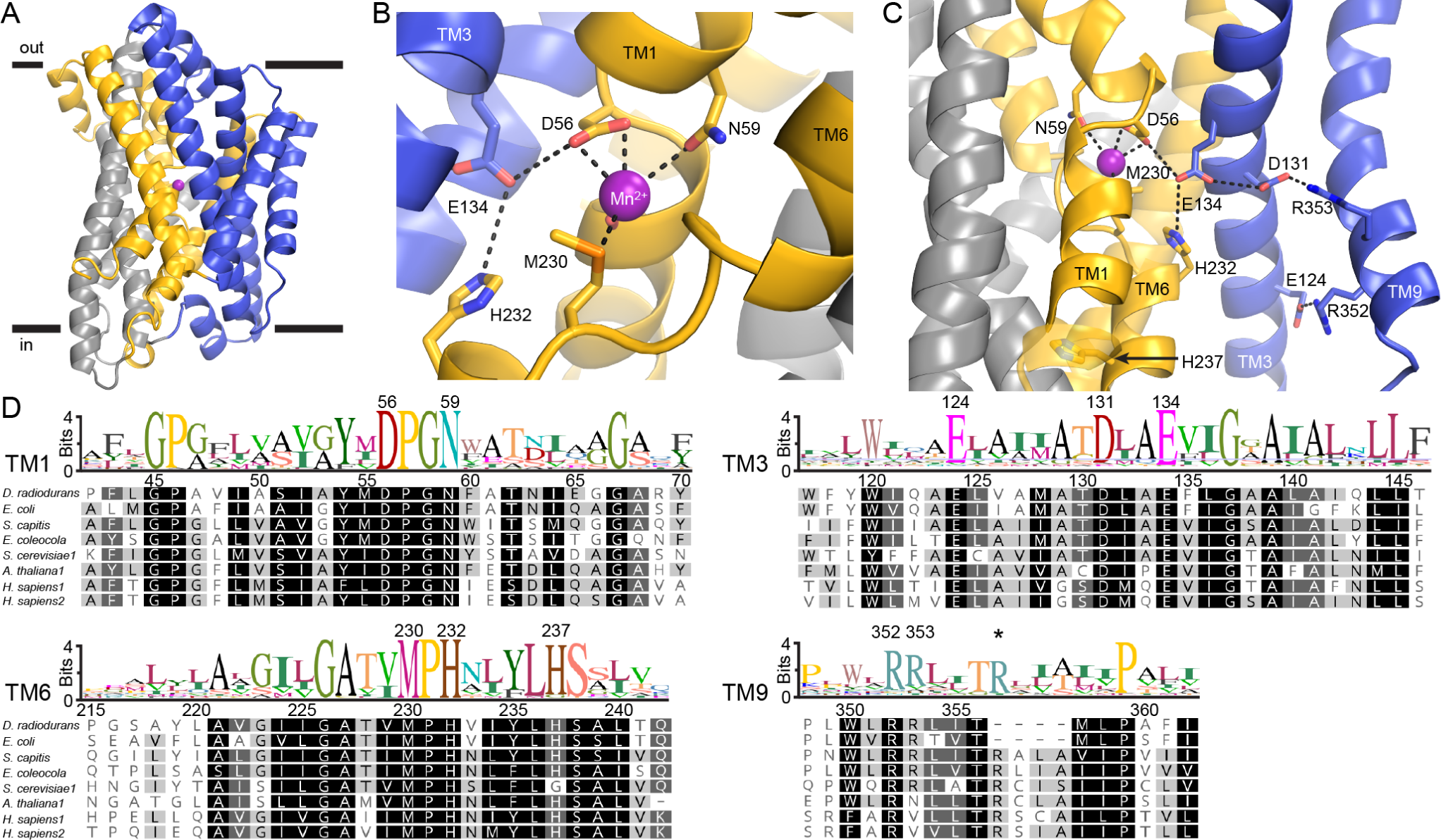
A network of highly conserved charged and protonatable residues adjoins the metal-binding site. (A) Crystal structure (PDB ID: 6BU5) of DraNramp in an outward-facing conformation bound to Mn^2+^ (magenta sphere). TMs 1, 5, 6, 10 are gold, TMs 2, 7, 11 gray, and TMs 3, 4, 8, 9 blue. (B) D56, N59, M230, and the A53 carbonyl coordinate Mn^2+^ in the outward-facing state, along with two waters (not shown). (C) View from the plane of the membrane of a network consisting of E134, H232, D131, R353, R352, and E124 that extends ∼20 Å from D56 to the cytoplasm. TMs 8 and 4, in front of TMs 3 and 9 respectively, were removed for clarity. (D) Sequence logos from an alignment of 6878 Nramp sequences. All 11 mutated residues (numbered above) are highly conserved. Canonical helix-breaking motifs at the metal-binding site are DPGN in TM1 and MPH in TM6. TM6’s H237 is a glycine in many fungal homologs. The second TM9 arginine (R353) varies in location (*) due to an extra helical turn in many homologs; this insertion contains a third arginine in some homologs. See also Figure 4—figure supplement 1.

In multiple Nramp homologs D56 and N59 contribute to metal transport (Bozzi et al., 2016a; Bozzi et al., 2018; Courville et al., 2008; Ehrnstorfer et al., 2014; Ehrnstorfer et al., 2017), while M230 contributes to a selectivity filter that favors transition metals over alkaline earth metals (Bozzi et al., 2016a). Mutations to E124, D131, E134, H232, and H237 abrogate Mn^2+^ transport in *E. coli* Nramp (Haemig and Brooker, 2004a; Haemig et al., 2010), while mutations to H232 and H237 (Lam-Yuk-Tseung et al., 2003; Mackenzie et al., 2006) and R353 (Lam-Yuk-Tseung et al., 2006) impair metal transport in human DMT1, with the latter mutation also causing anemia (Iolascon et al., 2006). Recent studies using different Nramp homologs implicated E134 or H232 in proton-metal coupling (Ehrnstorfer et al., 2017; Pujol-Gimenez et al., 2017), and we demonstrated the importance of D56 and D131 in addition to E134 and H232 to DraNramp H^+^-transport (Bozzi et al., 2018). Based on structural analyses we therefore proposed D56 as the initial proton-binding site and D131 as the subsequent proton acceptor required for transmembrane transport (Bozzi et al., 2018).

Hypothesizing that this network of conserved charged and protonatable residues contributes to the voltage-dependence and proton-metal coupling phenomena, we designed a panel of point mutants that remove or neutralize these sidechains via alanine and/or asparagine/glutamine substitution. All DraNramp mutants expressed well in *E. coli* (Figure 4—figure supplement 1C-D) and were readily purified. We reconstituted each in proteoliposomes to further explore how each conserved residue contributes to DraNramp’s metal and proton transport behavior.

### Conserved salt-bridge network imparts observed voltage dependence of metal transport

With our mutant panel, we first measured the effect of ΔΨ on *in vitro* Cd^2+^ and Mn^2+^ transport (Figure 5A and Figure 5—figure supplement 1). Most mutants retained some transport of either or both substrates, except metal-binding D56 mutants. Compared to WT, the remaining mutants cluster in two groups in terms of their ability to harness ΔΨ.

**Figure 5.**
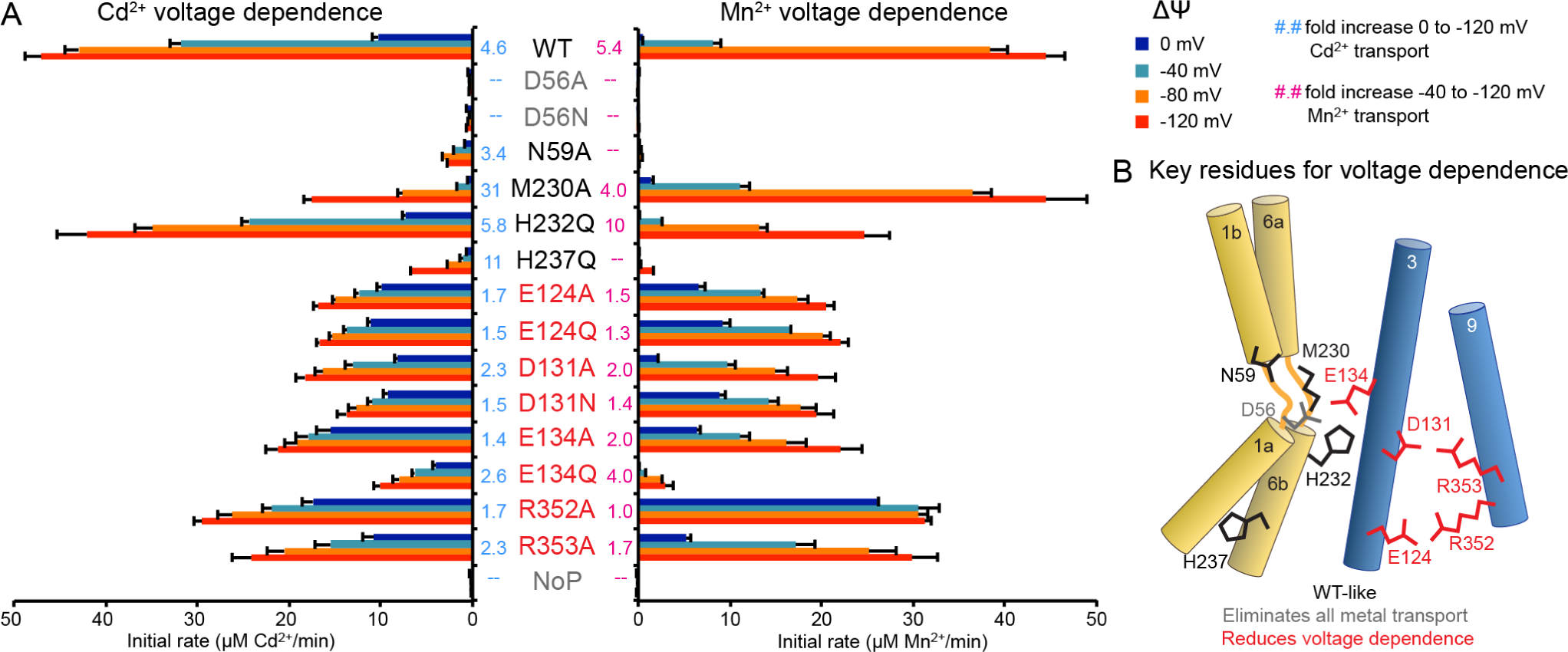
Conserved salt-bridge network confers observed voltage dependence. (A) Average initial metal uptake rates ± S.E.M. (n ≥ 4) for DraNramp mutants at different ΔΨ (colored bars), with the fold increase in transport indicated (cyan, 0 to −120 mV for Cd^2+^; pink, −40 to −120 mV for Mn^2+^). Mutations to E124, D131, E134, R352, and R353 (red) greatly reduced the ΔΨ dependence of Cd^2+^ and Mn^2+^ transport, such that they exceeded WT at low magnitude ΔΨ but lagged WT at physiological larger magnitude ΔΨ. Mutations to N59, M230, H232, and H237 (black) retained ΔΨ dependence, and D56 mutants (grey) eliminated all metal transport. (B) Network schematic illustrates clustering of residues most important for ΔΨ dependence (red). See also Figure 5—figure supplements 1 and 2.

Mutations to N59, M230, H232, and H237 largely preserved WT-like voltage dependence. These TM1 and TM6 residues cluster in the metal-binding site or metal-release pathway (Figure 5B). Mutations to TM3 and TM9 salt-bridge network residues E124, R352, R353, D131, and E134 reduced voltage dependence of Mn^2+^ and Cd^2+^ transport across the tested ΔΨs (Figure 5). All reduced transport rates at −120 mV, but many equaled and several outperformed WT at lower magnitude ΔΨ. Perturbing this network of residues thus alters ΔΨ’s effects on DraNramp’s metal transport kinetics, such that these mutants, while moderately impaired compared to WT under physiological conditions, no longer face WT’s transport restriction in the absence of ΔΨ.

We saw similar trends with this panel *in vivo*: salt-bridge network mutations reduced the effect of high external [K^+^] on Co^2+^ transport in *E. coli* (Figure 5—figure supplement 2), most strikingly with E134A and R352A. Overall these results further implicate the intact salt-bridge network as contributing to DraNramp’s strong voltage dependence and imposing a restriction on most metal transport when ΔΨ ≥ 0.

### Metal-binding site and salt-bridge network contribute to DraNramp’s proton-metal coupling

We next sought to disentangle the overlapping effects of ΔpH-stimulated metal transport and metal-stimulated proton co-transport, which we refer to as proton-metal coupling. We measured Mn^2+^ transport by our mutants in liposomes across a range of external pHs (and thus pH gradients) both in the presence and absence of a ΔΨ < 0 (Figure 6A and Figure 6—figure supplement 1A-B). Mutant phenotypes again clustered in three groups (Figure 6B). As before, mutants to metal-binding D56 and N59 eliminated Mn^2+^ transport, while H237Q showed slight activity with combined ΔΨ and ΔpH. As with voltage dependence, all mutants to the E124-R352-R353-D131-E134 network reduced the Mn^2+^-transport enhancement provided by ΔpH, to a level comparable to Cd^2+^ in WT. Strikingly, mutants M230A, H232Q, and E134Q—all near metal-binding D56— eliminated ΔpH stimulation, such that unlike the WT, Mn^2+^ transport was not observed in the absence of ΔΨ, no matter the ΔpH (Figure 6—figure supplement 1A-B).

**Figure 6.**
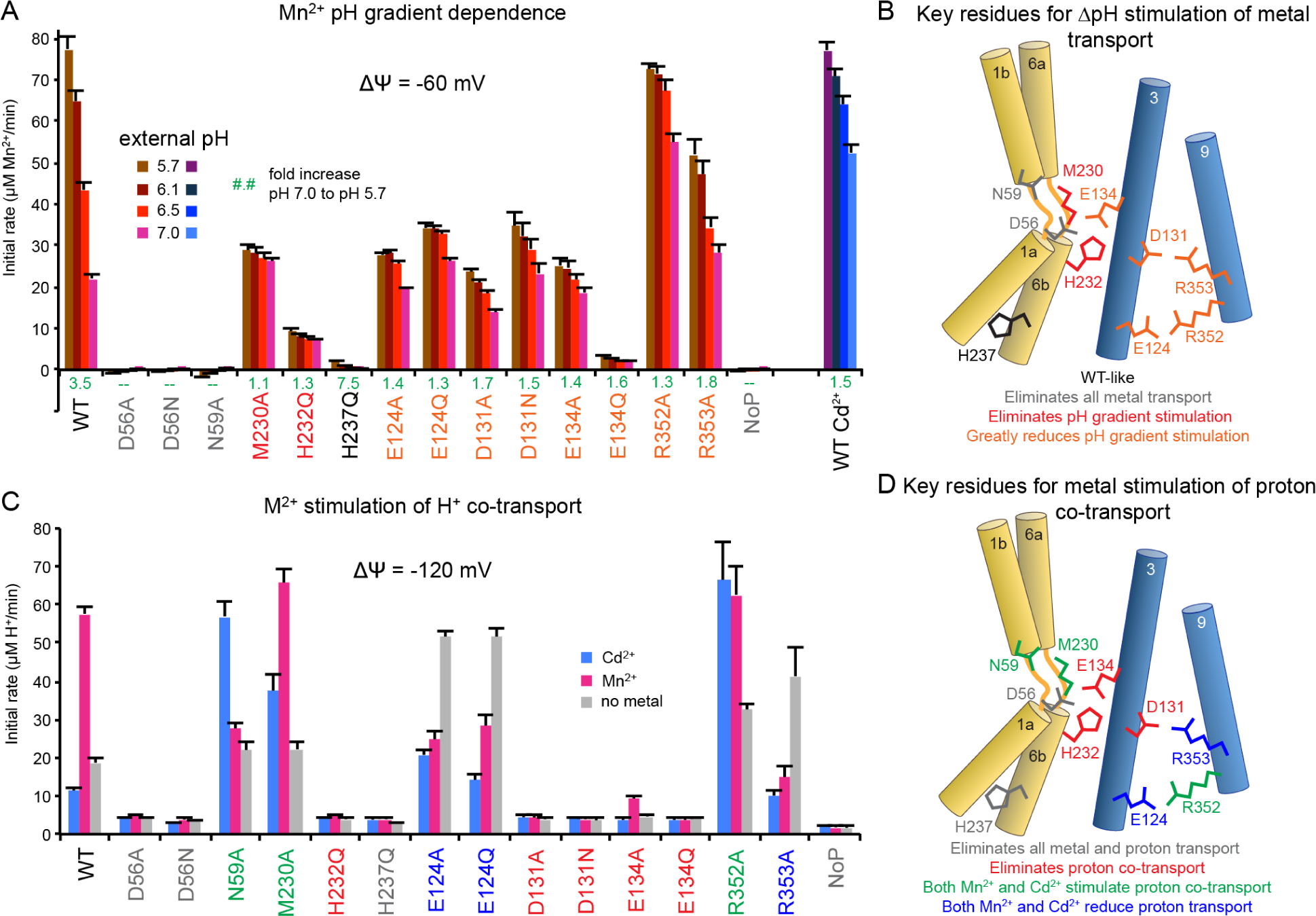
Entire conserved network enables ΔpH stimulation and enforces proton co-transport. (A) Average initial Mn^2+^ uptake rates ± S.E.M. (n ≥ 4) in proteoliposomes at various external pHs at ΔΨ = −60 mV. (B) Schematic shows proximity to binding site of essential residues for ΔpH stimulation (red), while entire extended network makes some contribution to ΔpH effect (orange). (C) Average initial metal-stimulated H^+^ transport rates ± S.E.M. (n ≥ 4). (D) Schematic shows that residues in immediate vicinity of D56 are essential for metal-stimulated H^+^ transport, while the entire network affects metal-proton coupling behavior. Interestingly, D131 and E134 mutants retain some ΔpH stimulation while eliminating metal-stimulated H^+^ co-transport, while M230A shows the opposite pattern. See also Figure 6—figure supplements 1 and 2.

Our mutants grouped differently when measuring how metal substrates Cd^2+^ and Mn^2+^ affect proton transport at −120 and −80 mV (Figure 6C-D, Figure 6—figure supplement 1C, and Figure 6—figure supplement 2). Mutations to D56, H232, E134, and D131 eliminated both basal and metal-induced H^+^ transport (E134A retained slight Mn^2+^ stimulation). In contrast to its inhibition of WT, Cd^2+^ stimulated H^+^ uptake for mutants to metal-binding N59 and M230 and the 18 Å-distant R352. Strikingly, for the E124 and R353 mutants, either Mn^2+^ or Cd^2+^ sharply reduced H^+^ transport—thus Mn^2+^ now behaved more like Cd^2+^ did for WT.

Overall, any mutation to the extended network perturbed proton-metal coupling and eliminated the distinction between Cd^2+^ and Mn^2+^. This is consistent with previous observations of subtle perturbations to this network affecting proton-metal coupling, as seen with an anemia-causing mutation G185R on TM4 in mouse Nramp2 (Fleming et al., 1998; Su et al., 1998) (G153R in DraNramp) directly above the R353-D131 pair (Xu et al., 2004), and an F-to-I substitution in human Nramp2 at the DraNramp equivalent L164 adjacent to R352-E124 (Nevo and Nelson, 2004). Curiously, some mutants (D131A, E134A) retained some ΔpH stimulation of Mn^2+^ transport but little Mn^2+^ stimulation of H^+^ transport, while others (M230A, R352A) had the opposite effect, with little ΔpH stimulation but robust Mn^2+^-stimulation. Interestingly, H232Q eliminated both effects—consistent with the H-to-A mutant in *Eremococcus coleocola* Nramp (Ehrnstorfer et al., 2017)—while D56 mutations eliminated all transport of metal and proton. These results indicate that pH stimulation of metal transport and metal-proton co-transport are distinct phenomena, likely imparted by separate groups of conserved residues.

### Perturbing the salt-bridge network impairs Mn^2+^ transport under physiological conditions

As high transport efficiency requires selectivity against potential competing substrates, we tested whether our mutants transport Ca^2+^, an abundant alkaline earth metal that Nramps must discriminate against (Figures 7A-B and Figure 7—figure supplement 1A-B). Most mutations did not increase Ca^2+^ transport. Thus, their perturbations of proton-metal coupling and voltage dependence do not arise from indiscriminate metal transport. Metal-binding D56 and N59 mutants reduced Ca^2+^ transport below WT levels, indicating that Ca^2+^ transport requires the same metal-binding site as other metals. Two mutations enhanced Ca^2+^ uptake: M230A, which removes the metal-coordinating methionine that preferentially stabilizes transition metals (Bozzi et al., 2016a), and R353A. The altered selectivity of R353A, 15 Å from the bound metal, is reminiscent of G153R, which adds exogenous positive charge to the salt-bridge network region and perturbs the metal-binding site to improve Ca^2+^ transport (Bozzi et al., 2016b; Xu et al., 2004).

**Figure 7.**
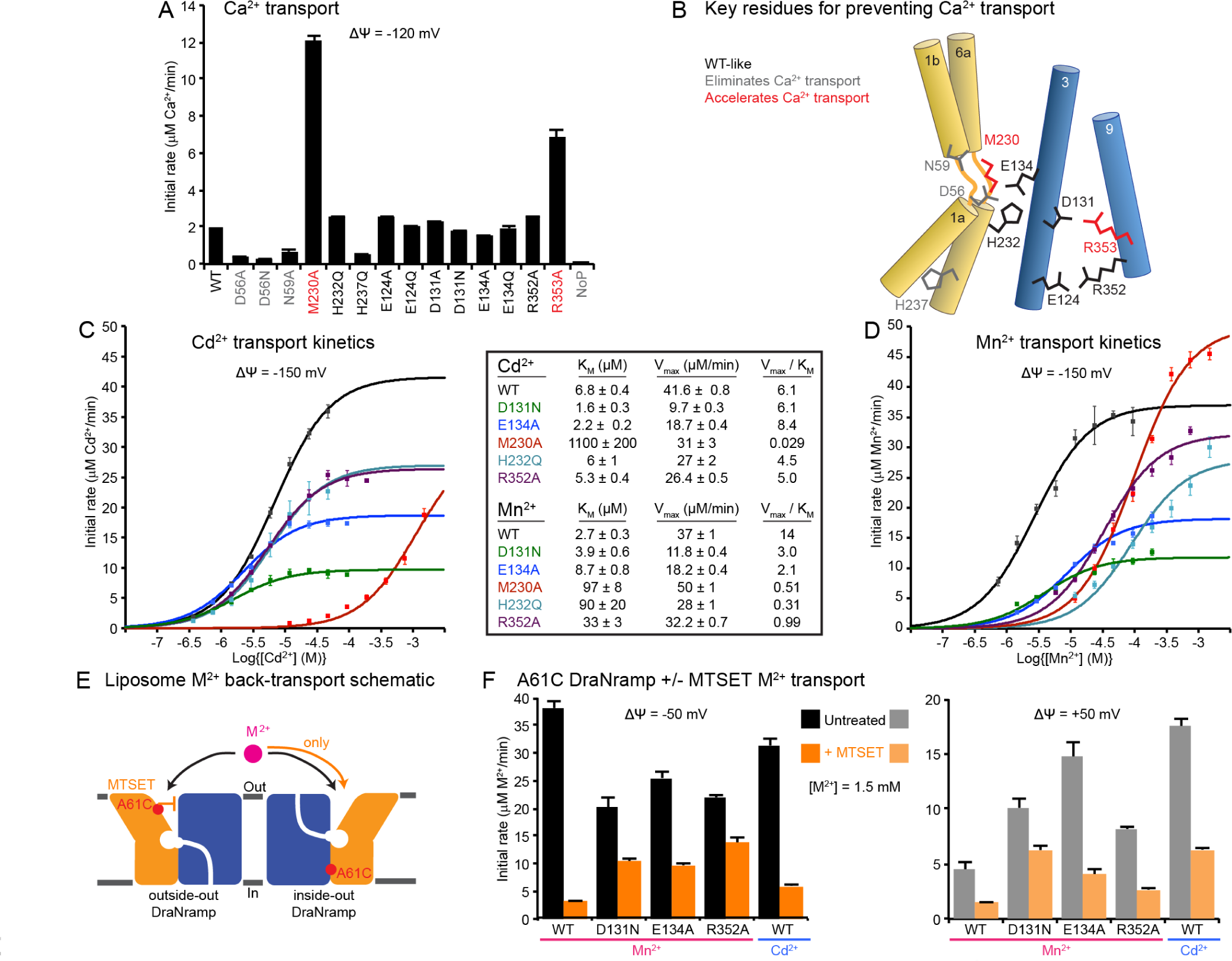
Perturbations to salt-bridge network impair high affinity Mn^2+^ uptake and enable deleterious Mn^2+^ back transport. (A) Average initial Ca^2+^ transport rates ± S.E.M. (n ≥ 4). (B) Schematic of mutation locations. M230A, or R353A, 15 Å away, both increased Ca^2+^ transport. D56 and N59 mutations eliminated Ca^2+^ uptake, and other mutants were similar to WT. (C-D). Dose-response curves of Cd^2+^ (C) or Mn^2+^ (D) transport for a subset of mutants that reduce or eliminate ΔΨ dependence, ΔpH stimulation, or proton co-transport. Data are averages ± S.E.M. (n ≥ 2). The resulting transport kinetic values (middle) show significant overlap for Cd^2+^ transport but wider separation for the physiological substrate Mn^2+^. M230A is the only mutation that impaired Cd^2+^ transport more severely than Mn^2+^ transport. Errors in KM and Vmax encompass the uncertainty of the fit to the data. (E) Schematic for isolating metal-transport activity from the ∼50% inside-out transporters. Only outside-out transporters are incapacitated by A61C modification with MTSET. (F) Average initial transport rates ± S.E.M. (n = 4) of MTSET-treated or untreated proteoliposomes. MTSET essentially eliminated WT-like’s Mn^2+^ transport. In contrast, D131N, E134A, and R352A retained significant Mn^2+^ transport, likely reflecting increased activity from inside-out transporters. This Mn^2+^ back transport still occurs under physiological-like conditions (ΔΨ > 0 for inside-to-outside transport). In addition, the WT-like protein retained greater residual transport of Cd^2+^, which did not require H^+^ symport (Figure 2). See also Figure 7—figure supplement 1.

We next compared Cd^2+^ and Mn^2+^ transport by select mutants across a range of metal concentrations to understand how these residues impact transport under more physiologically-relevant conditions (Figure 7C-D). Most mutations had little impact on Cd^2+^ transport even at low μM concentrations, and thus retained transport efficiency (calculated as Vmax/KM) within 2-fold of WT. The exception was M230A, which drastically reduced Cd^2+^ transport, consistent with the importance of this residue for Cd^2+^ affinity (Bozzi et al., 2016a; Bozzi et al., 2018). In contrast, all mutations reduced Mn^2+^ transport at environmentally-relevant low Mn^2+^ concentrations, resulting in 5 to 50-fold lower efficiency. Thus, mutagenic perturbations to the transporter that reduce voltage dependence or eliminate the proton co-transport requirement also impair function under physiological conditions. That these mutations alter DraNramp’s relative substrate preferences further supports an allosteric functional link between the salt-bridge network and the metal-binding site.

### Voltage dependence and proton-metal coupling prevent deleterious Mn^2+^ back-transport

Lastly, we probed whether the salt-bridge network prevents back-transport of metal ions. As membrane proteins typically orient randomly in proteoliposomes, we reconstituted DraNramp constructs containing the single-cysteine A61C that lines the external vestibule and applied membrane-impermeable 2-(Trimethylammonium)ethyl methanethiosulfonate (MTSET). Covalent modification of A61C with MTSET nearly eliminates metal transport *in vivo* (Bozzi et al., 2016b) and *in vitro* (Bozzi et al., 2018), likely by preventing outward-to-inward conformational change essential for metal transport (Bozzi et al., 2018) (Figure 7—figure supplement 1C). In proteoliposomes, MTSET treatment should thus incapacitate outside-out transporters, leaving inside-out proteins unaffected—and thus theoretically capable of metal transport (Figures 7E and S8D). Strikingly, while MTSET essentially eliminated Mn^2+^ transport for the WT-like protein, salt-bridge network mutants D131N, E134A, and R352A retained significant activity at both positive and negative ΔΨ (Figure 7F and Figure 7—figure supplement 1E). Thus inside-out mutant transporters shuttled Mn^2+^ down its concentration gradient against ΔΨ —corresponding to back-transport in a cellular context (Figure 7—figure supplement 1F)—while this process was only minimally observed for the WT-like protein. MTSET-treated WT-like protein facilitated greater Cd^2+^ transport, which unlike Mn^2+^ does not stimulate H^+^ uptake (Figure 2), further underscoring the importance of proton co-transport to enforce unidirectional metal transport. In conclusion, disruption of voltage dependence and proton-metal coupling through mutagenesis or by metal identity imparts a greater risk of metal efflux under physiological conditions.

## Discussion

We summarize the results of our mutagenesis experiments in Table S1, which inform the following model (Figure 8A). The D56-E134-D131-R353-R352-E124 network confers strong voltage dependence to the transporter, such that efficient metal transport only occurs at significantly negative ΔΨ (Figures 1 and 5). D56 first protonates to optimally orient the metal-binding residues—donating a hydrogen bond to the metal-interacting N59 carbonyl oxygen rather than receiving a hydrogen bond from the amide nitrogen (Figure 8B)—a process facilitated by E134, M230, and H232. D56 protonation also neutralizes the protein core. Incoming Mn^2+^ displaces the H^+^ from D56 and interacts with N59, the A53 carbonyl, and M230, which bonds semi-covalently to selectively stabilize the transition metal substrate. H232 and E134 may stabilize the H^+^ in a transition state before it reaches D131. This H^+^ transfer thus redistributes the net added positive charge, such that both the metal-binding site and the salt-bridge network gain a +1 formal charge. Next, the proton is released to the cytoplasm through the salt-bridge network. Lastly, bulk conformational change brings the transporter to the inward-facing state, in which the external vestibule closes and the cytosolic vestibule opens to allow eventual Mn^2+^ release (Bozzi et al., 2018). The salt-bridge network, a distinguishing feature of the Nramp clade of the LeuT family, thus amplifies voltage dependence and provides for temporal and spatial separation of the H^+^ and Mn^2+^ movement.

**Figure 8.**
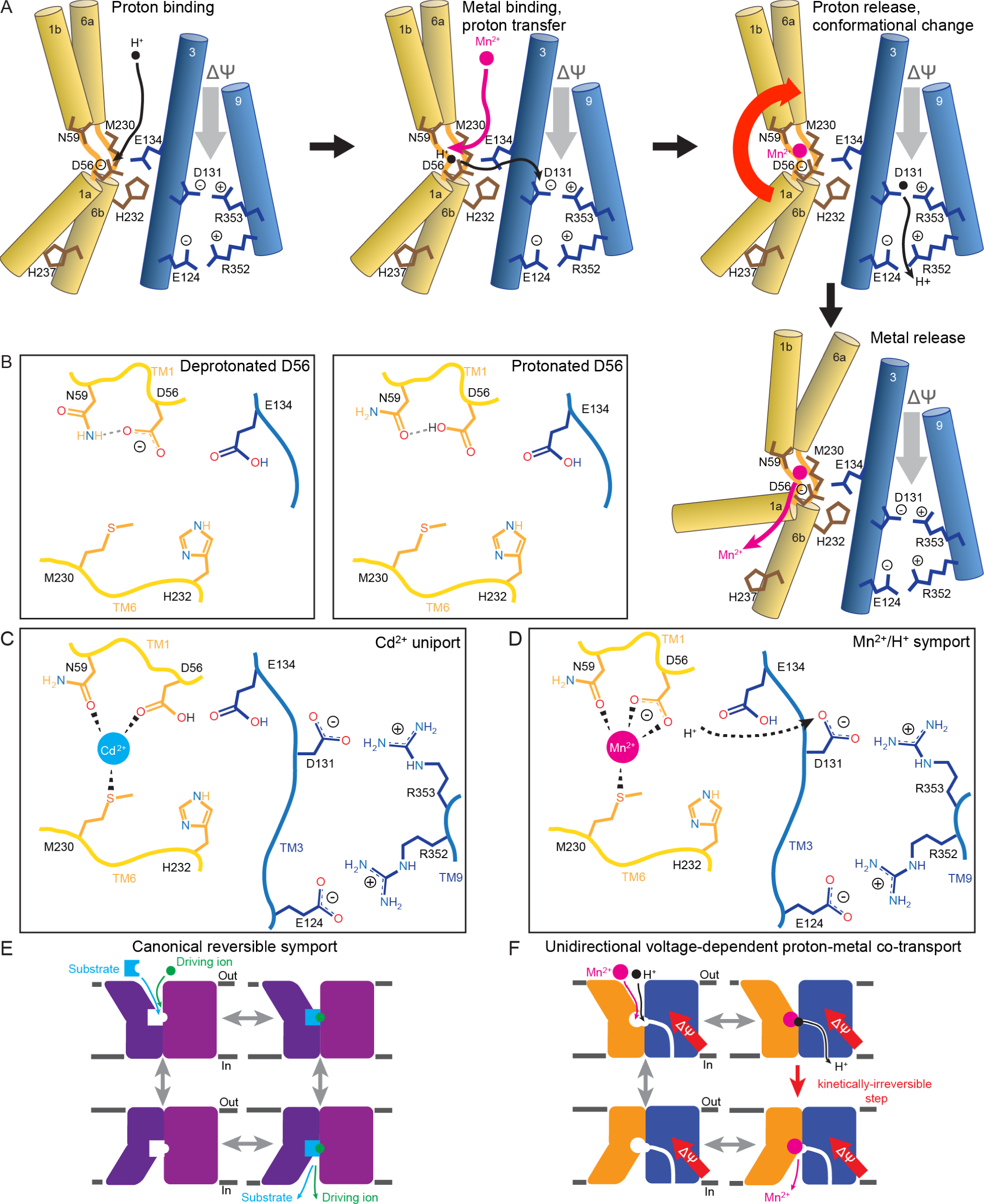
Complete mechanistic model for DraNramp transport processes. (A-B) Proposed mechanism of voltage-dependent, metal-proton symport. (Top left) D56 protonation optimizes the metal-binding site via hydrogen-bonding of D56 to the N59 carbonyl oxygen, providing a better metal-binding ligand than the amide nitrogen (as shown in B). (Top center) Metal substrate binds, displacing the D56 proton, which passes to D131, with H232 and E134 stabilizing a high-energy transition state. (Top right) The proton exits to the cytoplasm via the TM3/TM9 salt-bridge network. In tandem, a bulk conformational change closes the external vestibule and opens the cytoplasmic vestibule. (Bottom right) The metal is released to the cytoplasm. (C-D) Cd^2+^ uniport perhaps occurs due to a monodentate interaction with D56, enabling proton retention. Bidentate binding of D56 by Mn^2+^ requires deprotonation, passing the proton to D131. (E-F) Models for canonical secondary transport and DraNramp Mn^2+^/H^+^ transport. (E) In canonical secondary transporters, each step of the mechanism is reversible and tight coupling of the co-substrates ensures that the prevailing electrochemical gradients determine the net directionality of transport. (F) In contrast, co-substrate coupling is loose for DraNramp due to the requisite spatial separation of the like-charged Mn^2+^ and H^+^ during transport. In addition, the salt-bridge network that imparts voltage dependence and provides the proton-transport pathway enforces unidirectional Mn^2+^ transport, i.e. the Mn^2+^-bound inward-to-outward transition is kinetically blocked. See also Figure 8—figure supplement 1.

Substrate-specific differences in stoichiometry of the driving ion, as we demonstrated for DraNramp (Figure 2), have been reported for unrelated transporters (Chen et al., 1999b; Dohan et al., 2007), and different metal ions can trigger distinct allosteric responses upon binding, especially in metal-sensor proteins (Helmann, 2014; Waldron and Robinson, 2009). For DraNramp, the distinction between proton-metal symport (Mn^2+^, Fe^2+^, and Co^2+^) and metal uniport (Zn^2+^ and Cd^2+^) follows a familiar inorganic chemistry partition. The former cations have 5, 6, or 7 valence d-electrons and typically prefer a coordination arrangement with more electron-donating ligands than do the latter metals which have a filled d-shell (Barber-Zucker et al., 2017). Therefore, while D56 coordinates Mn^2+^ in a bidentate manner requiring deprotonation (Figure 8D) as we modeled for DraNramp’s outward-facing state (Bozzi et al., 2018), D56 may instead interact with Cd^2+^ only through its carbonyl oxygen and thus retain its proton (Figure 8C). The greater importance of M230 for Cd^2+^ than Mn^2+^ transport (Figure 7C-D and Bozzi et al., 2016a; Bozzi et al., 2018) and the necessity of a proposed transient metal-binding residue Q378 for high-affinity Mn^2+^ but not Cd^2+^ transport (Bozzi et al., 2018), further supports differential binding properties for the two metals. Our Mn^2+^/H^+^ symport results are consistent with *in vitro* findings with Mn^2+^ for *E. coleocola* Nramp (Ehrnstorfer et al., 2017). In contrast, the Cd^2+^ uniport we report contradicts *in vivo* results showing Cd^2+^ stimulates intracellular acidification in cells overexpressing *E. coli* Nramp (Courville et al., 2008)—which has the same conserved salt-bridge network as DraNramp (Figure 4). However, this study did not address the effects of Mn^2+^ or other metal substrates on intracellular pH. Furthermore, the inherent complexity of such *in vivo* experiments compared to an artificial proteoliposome means that the observed intracellular pH changes were perhaps not due to flux through *E. coli* Nramp.

Thermodynamically, proton-coupled transport promotes M^2+^ uptake under favorable ΔpH and ΔΨ conditions and thus enables a more substantial cytoplasmic vs. external Δ[Mn^2+^] than Δ[Cd^2+^]. The distinction in co-transport stoichiometry in DraNramp may therefore have evolved to prevent the excessive accumulation of toxic Cd^2+^ while preserving the ability to amass large stores (Daly et al., 2004) of an essential resource like Mn^2+^.

Intriguingly, perturbing either the metal-binding site or the adjoining salt-bridge network removes the H^+^ co-transport distinction between Cd^2+^ and Mn^2+^ (Figure 6C). Cd^2+^ stimulated H^+^ co-transport by the N59A and M230A variants, suggesting that D56 may compensate for the loss of a native metal ligand by interacting in a bidentate manner with Cd^2+^ and thus shedding a H^+^. In addition, mutations at residues ≥ 15 Å from bound M^2+^ also upset this distinction—the R352A variant symports H^+^ with both metals, while E124A and R353A shift to uniport of both metals — further underscoring the long-range influence of the salt-bridge network.

Overall, our experiments with different metals and point mutations indicate two distinct roles for protons in DraNramp’s mechanism: (i) ΔpH stimulation of metal transport, and (ii) metal stimulation of proton co-transport, as we highlight in Figures 2 and 6. Lower pH likely optimizes the metal-binding site via D56 protonation, and mutations to residues in the immediate vicinity of D56 (M230, E134, H232) completely eliminate pH dependence of transport (Figure 6A-B), perhaps by shifting D56’s pKa. Yet low pH still accelerates Cd^2+^ transport in WT, and Mn^2+^ transport in proton-pathway mutants like D131A, although less than in WT. Complementarily, proton co-transport is still observed for mutants such as M230A that eliminate the effect of ΔpH but do not compromise the conserved proton-transport pathway (Bozzi et al., 2018). Therefore, a ΔpH effect on the uptake of primary substrate should not be conflated with H^+^ co-transport and *vice versa*, whether in other Nramp homologs or unrelated secondary transporters.

The canonical model for secondary transport posits a cycle in which each transition is fully reversible, precise stoichiometry is maintained, and the prevailing ion gradients determine the directionality of net transport (Figure 8E). Effective symport thus requires tight coupling between the co-substrates, which is implemented in transporters by the driving ions either structurally enabling binding of the primary substrate or mechanistically selectively stabilizing a certain conformational state (Perez and Ziegler, 2013; Rudnick, 2013). Indeed, for the LeuT amino acid transporter—a structural homolog of DraNramp—substrate binding depends on two bound Na+ ions (Erlendsson et al., 2017): one Na^+^ stabilizes the outward-open conformation (Claxton et al., 2010; Tavoulari et al., 2016) while a second Na^+^ binds the transported amino acid’s carboxylate (Yamashita et al., 2005).

The DraNramp proton-metal symport mechanism (Figure 8F) deviates from basic principles of canonical symport. First, substrate coupling is asymmetric: Mn^2+^ transport requires H^+^ co-transport, to the extent that an Mn^2+^ gradient cannot drive Mn^2+^/H^+^ transport unless H^+^ entry alone is thermodynamically favorable (from ΔΨ and/or ΔpH, as seen in Figures 1 and 2). In contrast, while Mn^2+^ stimulates H^+^ transport, H^+^ uniport still occurs readily—and without requiring bulk conformational change (Bozzi et al., 2018)—resulting in the variable co-transport stoichiometries previously reported for yeast and human Nramp homologs (Chen et al., 1999a; Mackenzie et al., 2007; Mackenzie et al., 2006). These “futile cycles” are energetically wasteful, depleting ΔΨ and ΔpH without assisting in the uptake of the primary substrate. It is intriguing that evolution has retained such a thermodynamic cost as a general feature of the Nramp family, especially as simple tweaks near the salt-bridge network were shown to reduce H^+^ uniport without impairing metal uptake (Nevo and Nelson, 2004). Second, in our model, the two substrates do not coexist in the binding site throughout the transport process: Mn^2+^ binding to D56 evicts the pre-bound H^+^ into its distinct exit pathway. Indeed, our results suggest that H^+^ transport and M^2+^ binding may actually become competitive processes under some conditions, underscoring the co-substrate’s imperfect synergy (Figure 3).

In addition, our results demonstrate an essential kinetic role for the physiological membrane potential in DraNramp metal transport. While voltage necessarily contributes to the net thermodynamics of moving charged substrates across the membrane, we observe a strong voltage dependence of the entire free energy landscape (Figure 3), such that a sufficiently negative ΔΨ is prerequisite for metal binding and/or transport to occur. Although voltage dependence may be a diffuse property for a transport protein, the conserved salt-bridge network lining the ion transport routes in Nramps likely evolved to amplify the inherent voltage dependence of electrogenic transport and establish delicate mechanistic restrictions—including metal-specific proton co-transport—which our mutagenesis data support (Figures 5-6).

The key outcome of these deviations from canonical secondary transport—loose proton-metal coupling and strong voltage dependence—is to enforce the unidirectionality of DraNramp metal transport (Figure 8F). Indeed, no Mn^2+^ transport occurred in the absence of a negative ΔΨ (Figure 1; cations leaving the cell would experience a net positive ΔΨ). Furthermore inside-out transporters—even with a favorable ΔΨ—fail to efficiently import Mn^2+^ into liposomes down a large concentration gradient (Figure 7F). The inward-to-outward metal-bound transition is thus essentially forbidden in the WT protein under physiological conditions. Strikingly, point mutations to the salt-bridge network that perturb proton-metal coupling and voltage dependence can lift these restrictions, such that these protein variants behave more like directionally-unbiased uniporters. DraNramp’s unique structural adaptation—a conserved salt-bridge network that allosterically amplifies the voltage dependence of metal cation transport and provides a multistep, one-way exit pathway to the cytoplasm for the loosely-coupled proton co-substrate—therefore allows it to avoid the liabilities of reversibility inherent in the canonical model for symport. Similar properly-tuned voltage dependence and asymmetric substrate coupling may be a more general strategy for transporters to prevent deleterious back-transport without jeopardizing transport in the desired direction.

## Methods

### In vivo Co^2+^ transport

Co^2+^ transport in *E. coli* was performed as described with 200 μM Co(NO3)_2_ (Bozzi et al., 2018), except that the assay buffer was 50 mM HEPES pH 7.0, 60 mM NaCl, 10 mM KCl, 0.5 mM MgCl_2_, 0.216% glucose. For the varied [K^+^] measurements, the assay buffer was 50 mM HEPES pH 7.0, 60 mM NaCl, 0.5 mM MgCl_2_, X mM KCl, and (82.5-X) mM choline chloride, with X indicated in the corresponding figure legends. For each biological replicate reported in the figure legends, a separate culture of transformed *E. coli* was grown and induced to express the exogenous Nramp construct.

### Cysteine accessibility measurements

Cysteine accessibility measurements in *E. coli* were performed as described (Bozzi et al., 2018), except that the assay buffer used was the same as for the Co^2+^ transport experiments. For each biological replicate reported in the figure legends, a separate culture of transformed *E. coli* was grown and induced to express the exogenous Nramp construct.

### Cloning, expression and purification of DraNramp constructs for proteoliposome assays

DraNramp constructs were cloned, expressed, and purified as described (Bozzi et al., 2018), with the following changes: protein was purified from cell pellets in a single day, and washed/eluted from nickel beads in buffers with 0.03% DDM. Protein was concentrated to 2.5 mL and buffer-exchanged into 100 mM NaCl, 10 mM HEPES pH 7.5, 0.1% n-Decyl-β-D-Maltopyranoside (DM) on a PD10 desalting column. Protein concentrations were normalized to 1.2 mg/ml and aliquots were flash frozen in liquid nitrogen. Single-cysteine constructs A53C and A61C were purified in the presence of 1 mM DTT.

### Proteoliposome preparation

Adjusting the lipid composition (Ehrnstorfer et al., 2017) of a previous protocol (Bozzi et al., 2016a; Tsai et al., 2014), 75% w/w 1-palmitoyl-2-oleoyl-sn-glycero-3-phosphoethanolamine (POPE) was mixed with 25% w/w 1-palmitoyl-2-oleoyl-sn-glycero-3-phosphoglycerol (POPG) in chloroform (Avanti Polar Lipids), and then dried under nitrogen in a warm water bath, re-dissolved in pentane, and dried again. Lipids were resuspended at 20 mg/ml in 5 mM DM in KCl+NaCl/MOPS buffer (typically ∼90 mM KCl, 30 mM NaCl, 0.5 mM or 10 mM MOPS pH 7). Protein was added at a 1:400 w/w ratio to lipid, and the mixture dialyzed at 4°C to remove the detergent in 10 kDa molecular weight cutoff dialysis cassettes against KCl+NaCl/MOPS buffer with 0.2 mM EDTA for 1 day, then with 0.1 mM EDTA for 1-3 days, then overnight at room temperature (RT) against KCl+NaCl/MOPS buffer. For A53C and A61C, 1 mM, and 0.5 mM DTT was included in the first two dialysis steps. Fluorescent dye (either 1:49 v/v 5 mM Fura-2 pentapotassium salt or 1:66 v/v 10 mM 2′,7′-bis(carboxyethyl)-5(6)-Carboxyfluorescein (BCECF) in dimethyl sulfoxide) was incorporated into proteoliposomes permeabilized by three freeze-thaw cycles in dry ice-ethanol and RT water baths (and sometimes stored at −80°C after the third freeze). Proteoliposomes were extruded through a 400 nM filter to create uniform-sized vesicles, buffer-exchanged 1-2 times on a PD10 desalting column pre-equilibrated with NaCl/ or KCl/10 mM MOPS pH 7 buffer. Peak proteoliposome-containing fractions were pooled to remove unincorporated dye.

### Proteoliposome transport assays and data analysis

Proteoliposomes loaded with either 100 μM Fura-2 or 150 μM BCECF were diluted into buffer containing appropriate [KCl] to establish the desired membrane potential(Fitzgerald et al., 2017; Uzdavinys et al., 2017) and aliquoted into 96 well black clear-bottom plates. Following baseline fluorescence measurement, 5X metal (750 μM final concentration unless otherwise noted) and valinomycin (100 nM final concentration) were added. A fresh stock of 100 mM FeSO_4_ in 100 mM L-ascorbic acid was made for each assay; stocks of 100 mM CdCl_2_, MnCl_2_, ZnCl_2_, and Co(NO_3_)_2_ were freshly diluted into appropriate NaCl or KCl buffer. For assays with a pH gradient, the metals were diluted into 100 mM MES at pH 5.5, 5.8, 6.0, or 6.5 with appropriate NaCl/KCl. The reported external pH upon metal addition was determined from proportional mixings of larger volumes of the same buffers. To pre-modify A61C constructs for transport assays, liposomes were diluted into 120 mM NaCl, 10 mM MOPS pH 7 containing 3 mM MTSET and incubated at least 30 min at RT before beginning transport assays.

Metal transport was monitored by measuring Fura-2 fluorescence at λ_ex_ = 340 and 380 nm, at λ_em_ = 510 nm. Proton transport was monitored by measuring BCECF fluorescence at λ_ex_ = 450 and 490 nm, at λ_em_ = 535 nm. To calculate concentrations of imported metal, the Fura-2 340/380 ratio and experimentally determined K_D_ values (Grynkiewicz et al., 1985; Hinkle et al., 1992) were used for Ca^2+^ and Cd^2+^ as described previously (Bozzi et al., 2016a); the Ca^2+^ K_D_ value was used as an approximation for Zn^2+^. For Mn^2+^, Fe^2+^, and Co^2+^, the fraction of Fura-2 340 and/or 380 fluorescence quenched, normalized to maximum observed quenching, was used to estimate imported metal. For proton uptake, the BCECF 450/490 ratio was used to calculate internal pH, which along with the known total internal buffer (0.5 mM) and dye (150 μM) concentration was used to calculate net proton import via the Henderson-Hasselbalch equation. Initial rates were calculated in Excel and Michaelis-Menten parameters were fit using MATLAB. For each technical replicate reported in figure legends, a separate aliquot of dye-loaded proteoliposomes was diluted into the appropriate outside buffer, including cysteine modifiers if applicable, then fluorescence time course data were collected before and after the addition of valinomycin, metal substrate, and/or ionomycin.

## Data availability

The raw biochemical data that support the findings of this study are available from the corresponding author upon reasonable request.

## Author contributions

R.G. oversaw and designed the research with A.T.B., L.B.B., and C.M.Z.; L.B.B. and A.T.B. developed the proteoliposome transport assay and performed preliminary experiments; C.M.Z. provided essential structural insights into the DraNramp transport mechanism that guided mutagenesis experiments; A.T.B. performed all experiments and analyzed the resulting data; A.T.B. and R.G. wrote the manuscript, with input from all authors.

## Acknowledgements

We would like to thank Sarah Farron for assistance with transport kinetics analysis, Casey Zhang for generating the Nramp sequence alignment, Chris Miller and Joe Mindell for critical feedback, and Niels Bradshaw, Jack Nicoludis, and the rest of the Gaudet lab for helpful discussions. This work was funded in part by NIH grant 1R01GM120996 (to R.G.) and a Jane Coffin Childs Postdoctoral Fellowship (to C.M.Z.).

---

**Figure 1—figure supplement 1.**
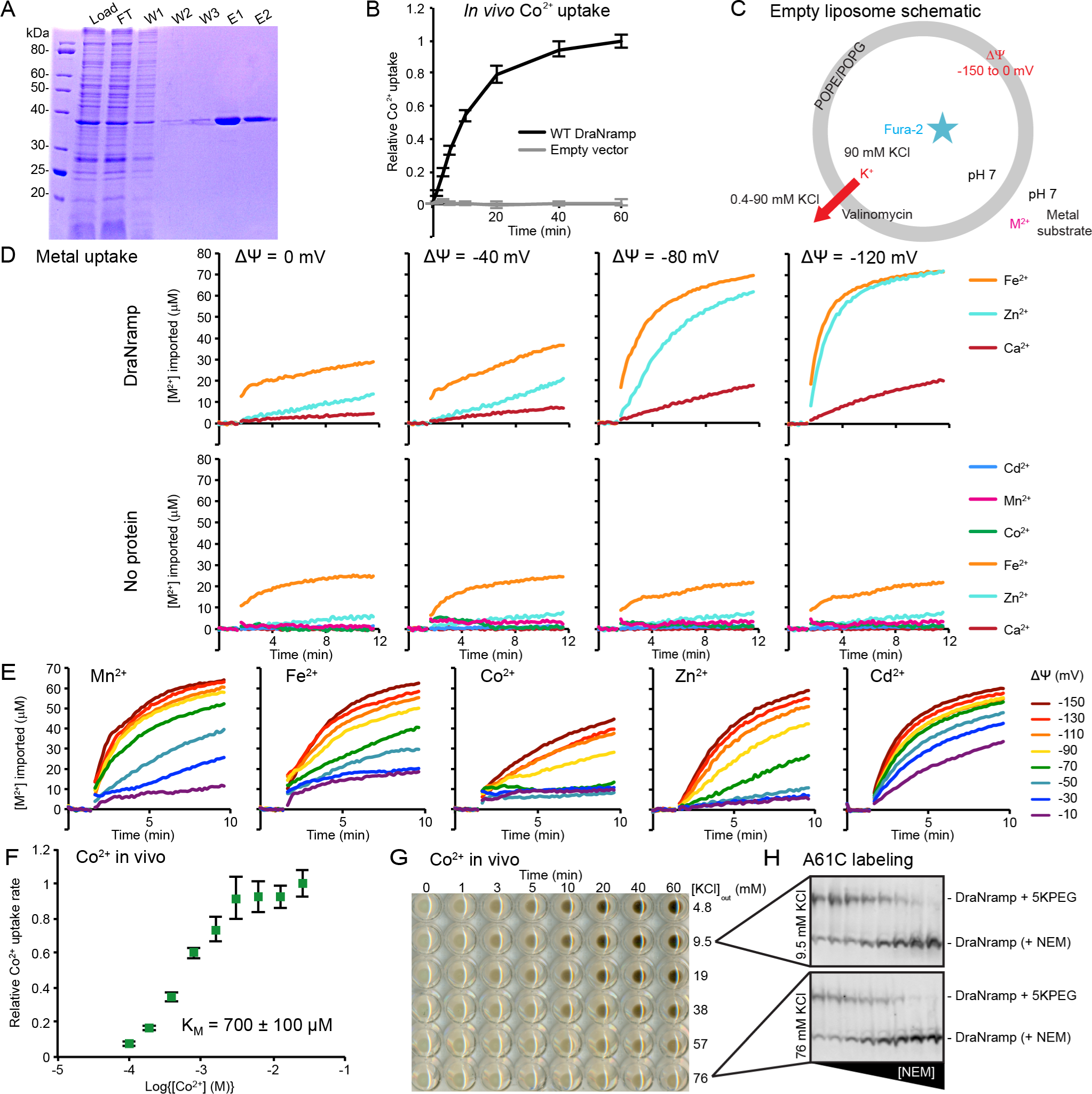
DraNramp is a voltage-dependent transition metal transporter. (A) Coomassie-stained SDS-gel of His-tagged DraNramp purification via Ni^2+^-affinity showing load, flowthrough (FT), washes (W1, W2, W3) and elutions (E1, E2). (B) Colorimetric detection of Co^2+^ taken up by cells over time shows that DraNramp transports Co^2+^ when expressed in *E. coli*. Data are averages ± S.E.M. (n = 4). (C) Control liposomes lacking DraNramp are used to determine protein-dependent transport activity. (D) Representative time traces (n ≥ 4) of metal uptake by DraNramp-containing liposomes (top) show that Fe^2+^ and Zn^2+^ also exhibit strong voltage dependence, while Ca^2+^ remains a poor substrate across all voltages. Control liposomes (bottom) have slow but non-negligible, voltage-independent Fe^2+^ and Zn^2+^ leak. (E) Representative time traces (n = 4) of metal uptake by DraNramp-containing liposomes at different membrane voltages. Transport of all five tested substrates is accelerated at high magnitude negative membrane potentials. (F) Dose-response curve shows that Co^2+^ transport in *E. coli* has K_M_ = 700 ± 100 μM similar to that obtained at ΔΨ = −150 mV in proteoliposomes, confirming that DraNramp behavior *in vitro* is physiologically relevant. Data are averages ± S.E.M. (n = 4). Error in K_M_ is the uncertainty of fit to data. (G) Representative image of Co^2+^ uptake assay results showing that increased extracellular [K^+^] inhibits DraNramp-dependent Co^2+^ uptake in *E. coli*, as detected by precipitating transported Co^2+^ as the dark solid CoS. (H) Representative western blots showing that A61C is labeled by N-ethylmaleimide (NEM) similarly in high or low external [K^+^].

**Figure 2—figure supplement 1.**
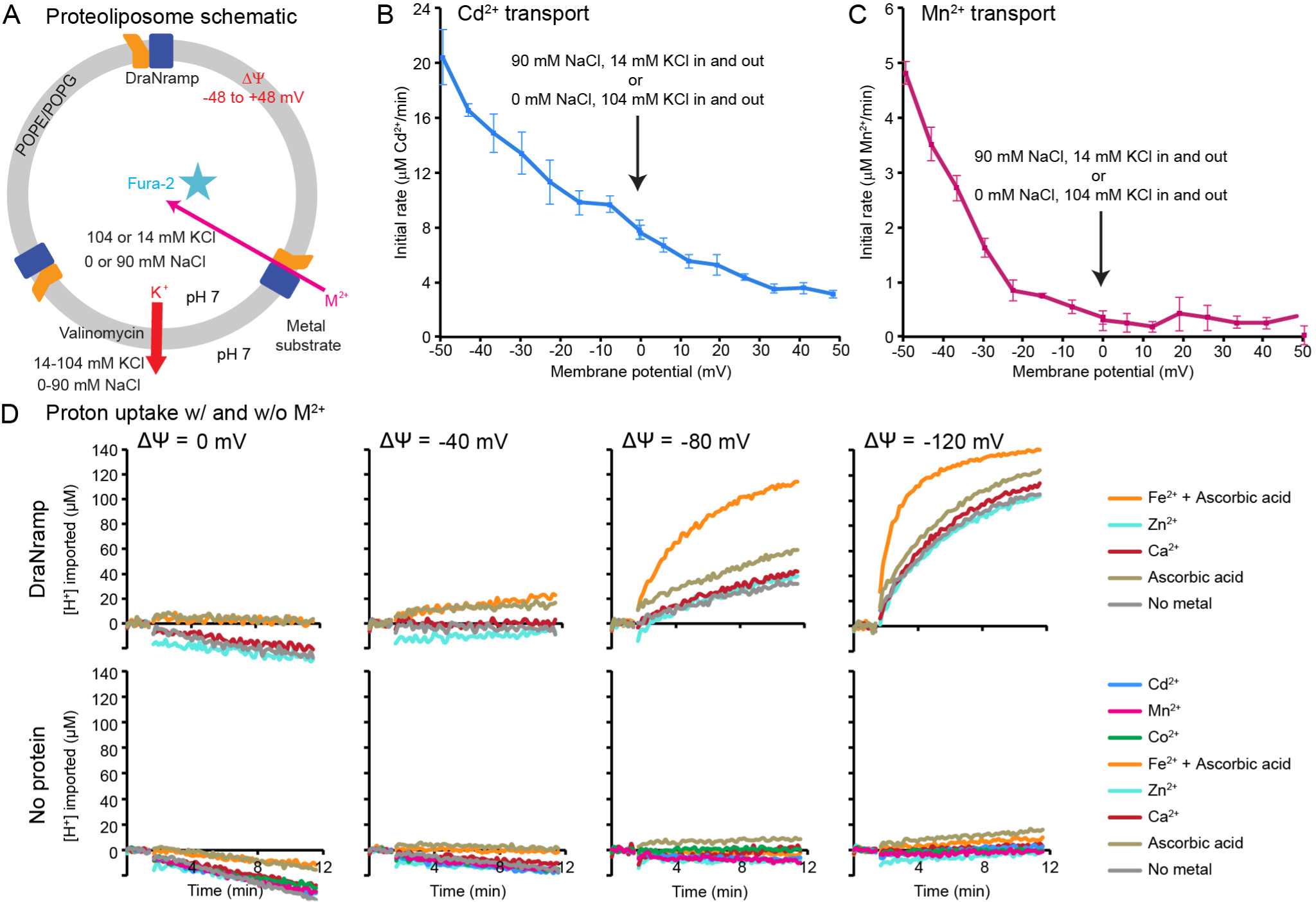
Proton co-transport is metal-specific. (A) Schematic for proteoliposome assays with moderate negative and positive ΔΨ values. Liposomes for ΔΨ ≤ 0 contained only KCl inside and were diluted to a lower external [K^+^], with increased Na^+^ used to balance osmolarity, before adding valinomycin to establish the membrane potential. Liposomes for ΔΨ ≥ 0 contained mainly NaCl inside, and were diluted into higher outside [K^+^]. (B-C) Average initial transport rate ± S.E.M. (n = 4) versus membrane potential. Cd^2+^ transport (B) driven by a [Cd^2+^] gradient occurs even at ΔΨ ≥ 0, although the rate decreases at more positive ΔΨ. In contrast, Mn^2+^ transport (C) does not occur unless ΔΨ < 0. Importantly, for both Cd^2+^ and Mn^2+^ at 0 mV, the observed transport behavior is the same regardless of whether the dominant cation species on both sides of the membrane is K^+^ or Na^+^. Thus, these ions on their own do not enable or inhibit transport, but rather the membrane potential causes the observed variations in transport rates. (D) Representative time traces (n ≥ 4) of H^+^ uptake. With proteoliposomes (top), Fe^2+^ stimulated H^+^ entry, while Zn^2+^ (substrate) and Ca^2+^ (non-substrate) did not. ΔΨ-driven H^+^ import was DraNramp-dependent, as there was no significant H^+^ leak or metal-stimulated H^+^ entry into control liposomes (bottom). Ascorbic acid used to prevent Fe^2+^ oxidation slightly lowered the external pH, which led to slight proteoliposome acidification. Except for the reporter dye, conditions were identical to those for the metal transport in Figure 1— figure supplement 1D.

**Figure 4—figure supplement 1.**
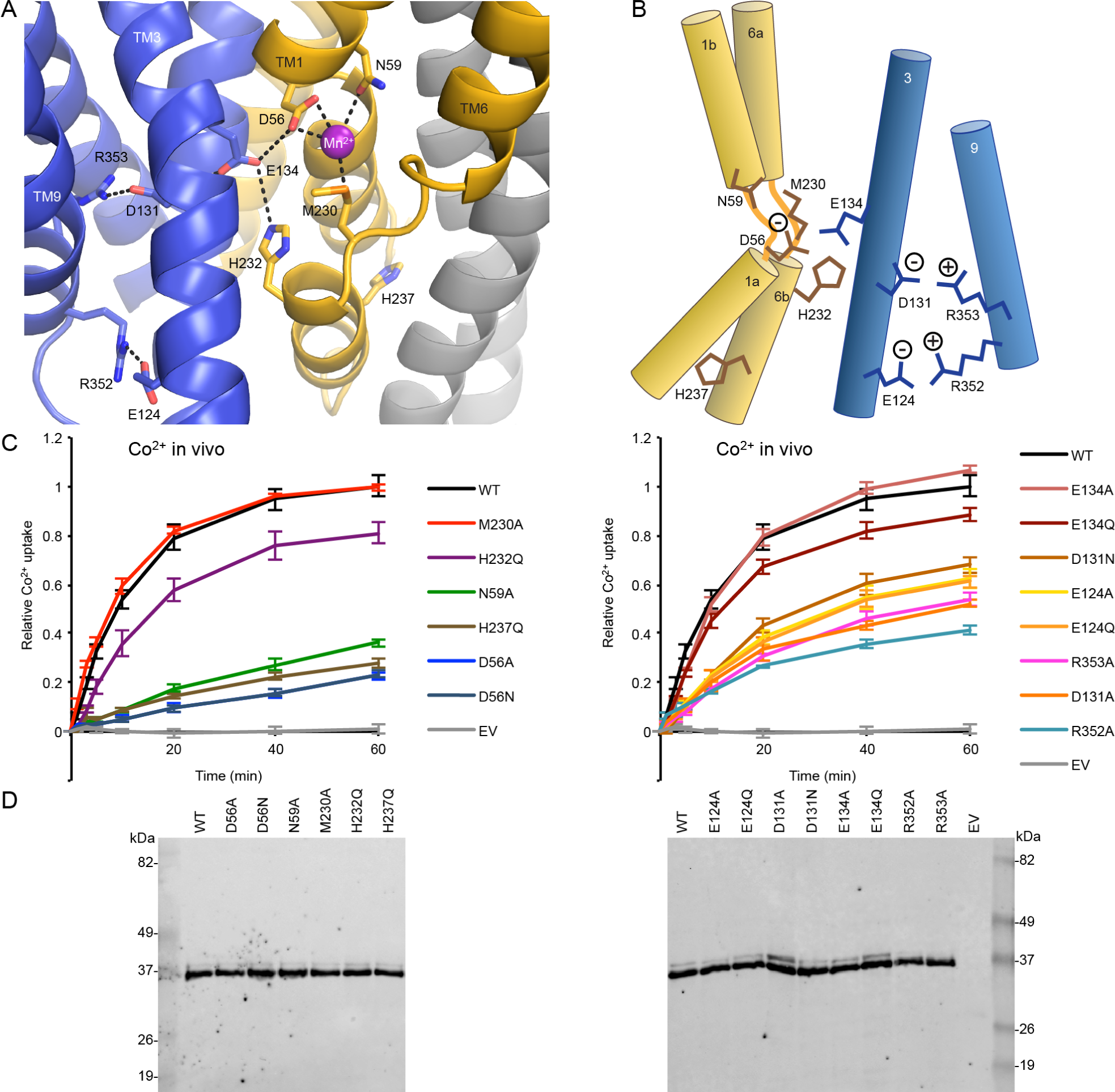
Conserved charged and protonatable residues form network leading from metal-binding site to the cytoplasm. (A) View down the metal entry pathway to the conserved network. TMs 10 and 11, which abut TMs 6a and 3, have been removed for clarity. TM6b’s H237 lines the metal exit pathway to the cytoplasm, while the rest of the network lines the proton exit route (Bozzi et al., 2018). (B) Schematic of the conserved residues in the salt-bridge network. (C) Time courses of *in vivo* Co^2+^ uptake (averages ± S.E.M.; n = 4). Mutants at most positions retained some Co^2+^ transport in *E. coli*, with D56, N59, and H237 mutants most impaired. (D) All constructs express similarly when detected via western blot for the N-terminal His-tag.

**Figure 5—figure supplement 1.**
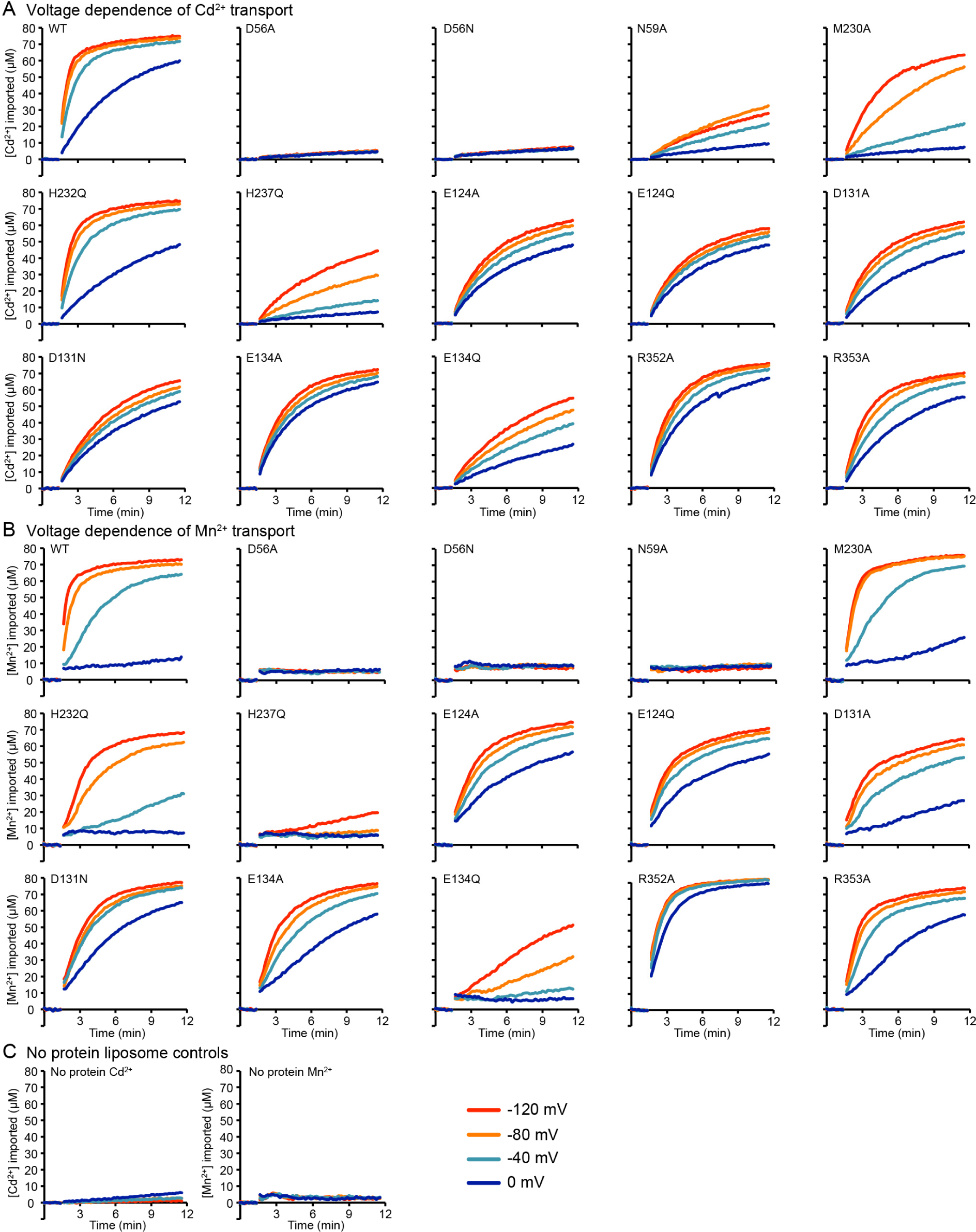
Sample traces illustrate mutant perturbations to voltage dependence. (A-B) Representative time traces (n ≥ 4) of Cd^2+^ (A) and Mn^2+^ (B) uptake into proteoliposomes measured at four ΔΨ values for each mutant. (C) Representative time traces (n > 4) of Cd^2+^ (left) and Mn^2+^ (right) uptake showed no metal was imported into control liposomes.

**Figure 5—figure supplement 2.**
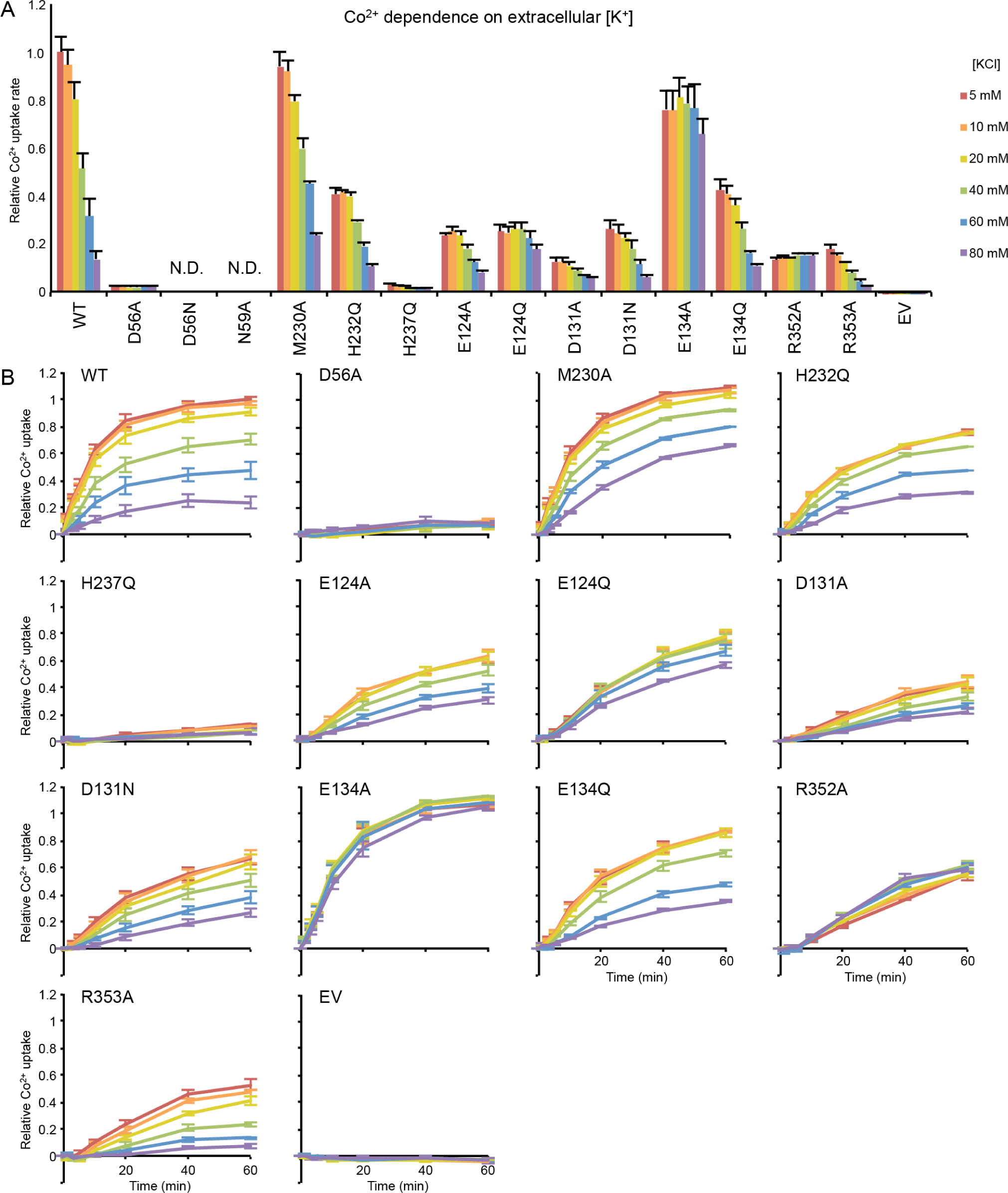
Mutations to conserved residues perturb voltage dependence of. ***in vivo* Co^2+^ transport.** (A) Relative Co^2+^ transport rates at various extracellular [K^+^] applied to perturb the endogenous *E. coli* membrane potential; higher extracellular [K^+^] presumably leads to a less-negative ΔΨ. Mutations to E124, D131, E134, and R352 reduced the effect of extracellular [K^+^] more than did mutations to M230, H232, and R353. D56N and N59A mutants were not tested. (B) Time course data used to generate the initial rates shown in (A). All data are averages ± S.E.M. (n ≥ 3).

**Figure 6—figure supplement 1.**
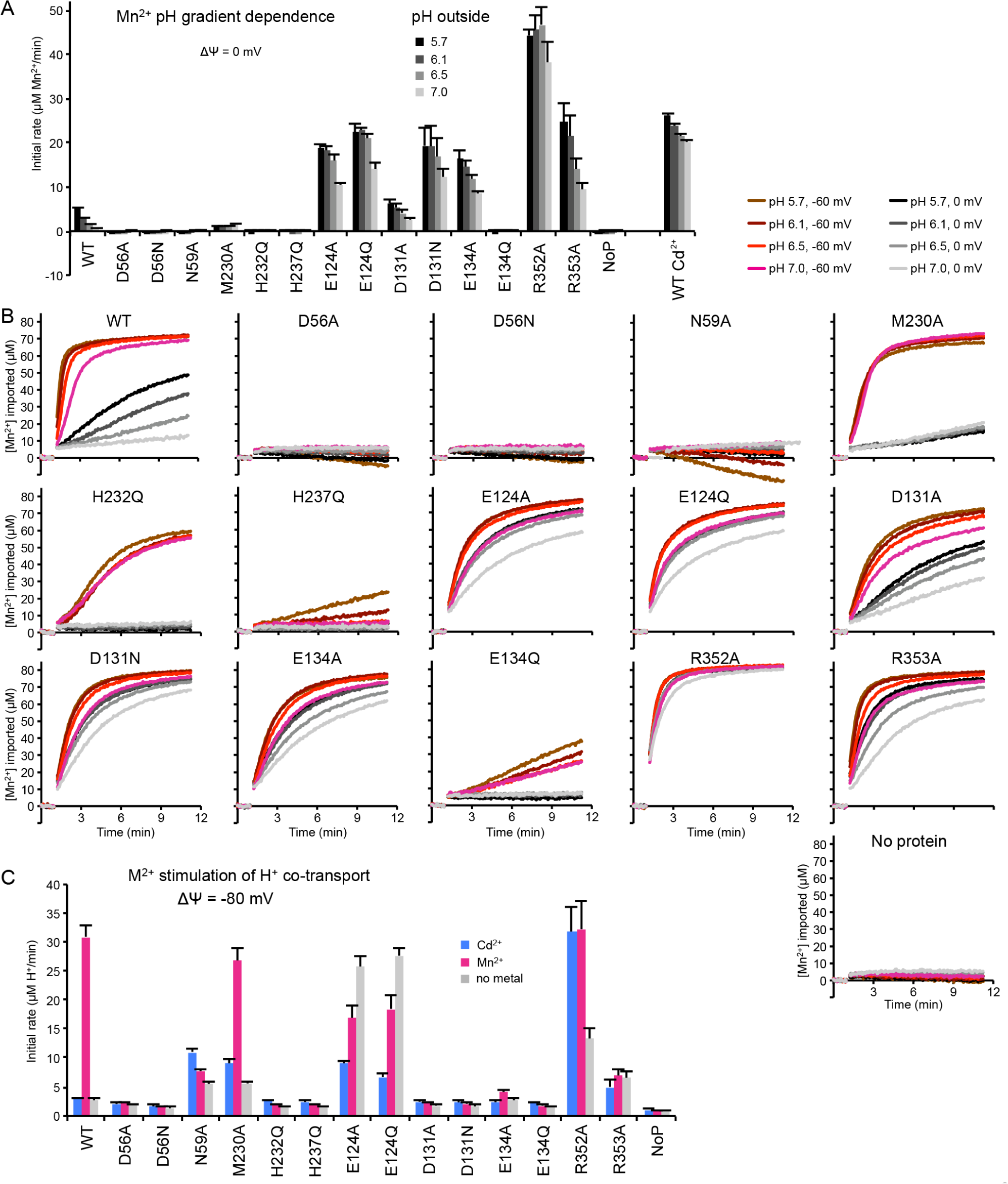
Effects of mutations on ΔpH and ΔΨ stimulation of Mn^2+^ transport. (A) Average initial Mn^2+^ transport rates ± S.E.M. (n ≥ 4) at various external pHs at ΔΨ = 0. E124, D131, E134, R352, and R353 mutants retained significant Mn^2+^ transport, but little ΔpH dependence. Even with a ΔpH, H232Q and M230A did not transport Mn^2+^. (B) Representative time traces (n ≥ 4) of Mn^2+^ influx at four ΔpH values at ΔΨ = 0 and −60 mV for each mutant. (C) Average initial metal-stimulated H^+^ transport rates ± S.E.M. (n ≥ 4) at −80 mV show the same trends as at −120 mV in Figure 6C.

**Figure 6—figure supplement 2.**
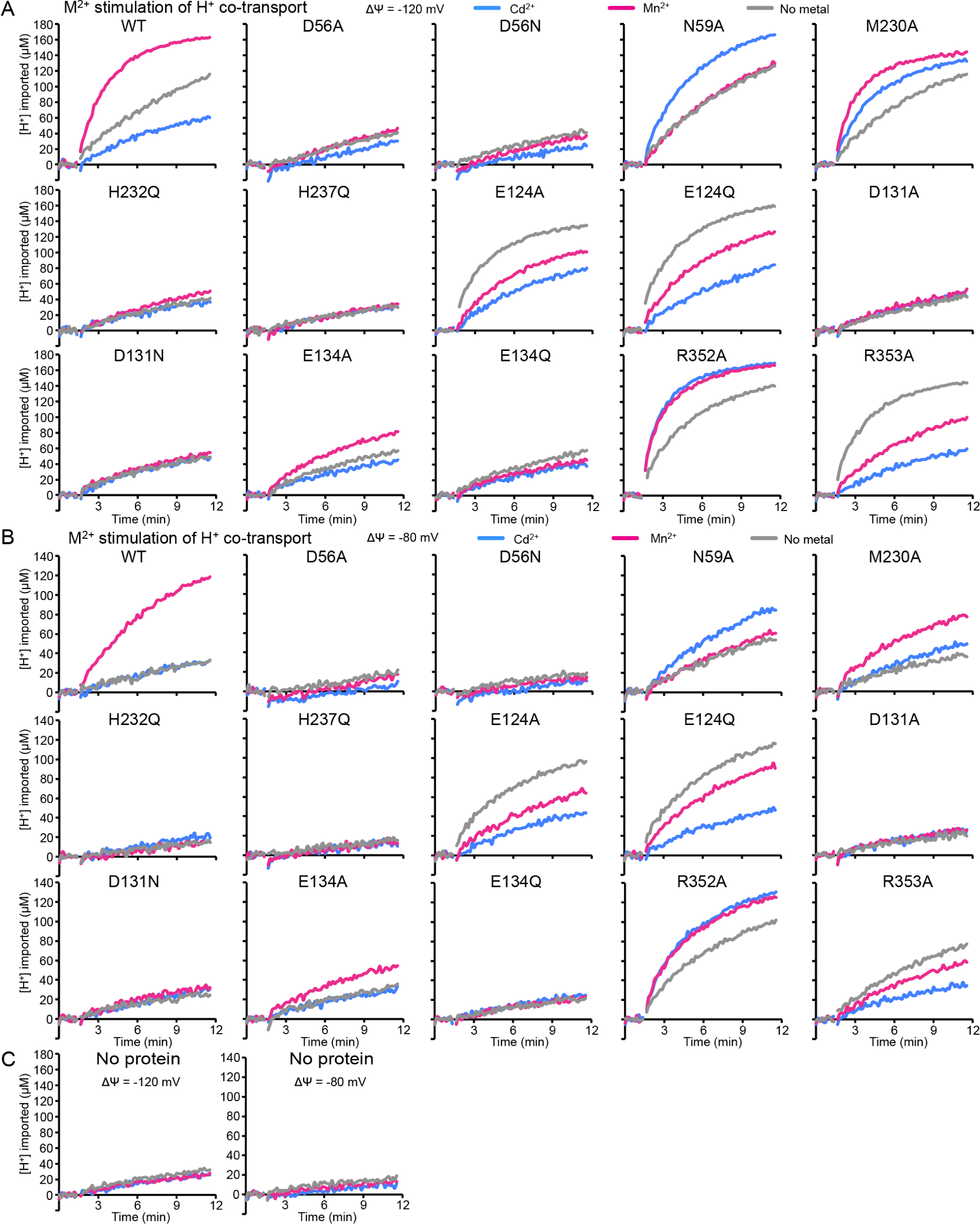
Sample traces show effects of mutations on metal-stimulated H^+^ co-transport. (A-B) Representative time traces (n ≥ 4) of metal-stimulated H^+^ transport in proteoliposomes at ΔΨ = −120 mV (A) or −80 mV (B). H^+^ entry was measured in the absence of metal and in the presence of Mn^2+^ or Cd^2+^ for each mutant. (C) No significant H^+^ entry was observed into control no protein liposomes under any of the tested conditions.

**Figure 7—figure supplement 1.**
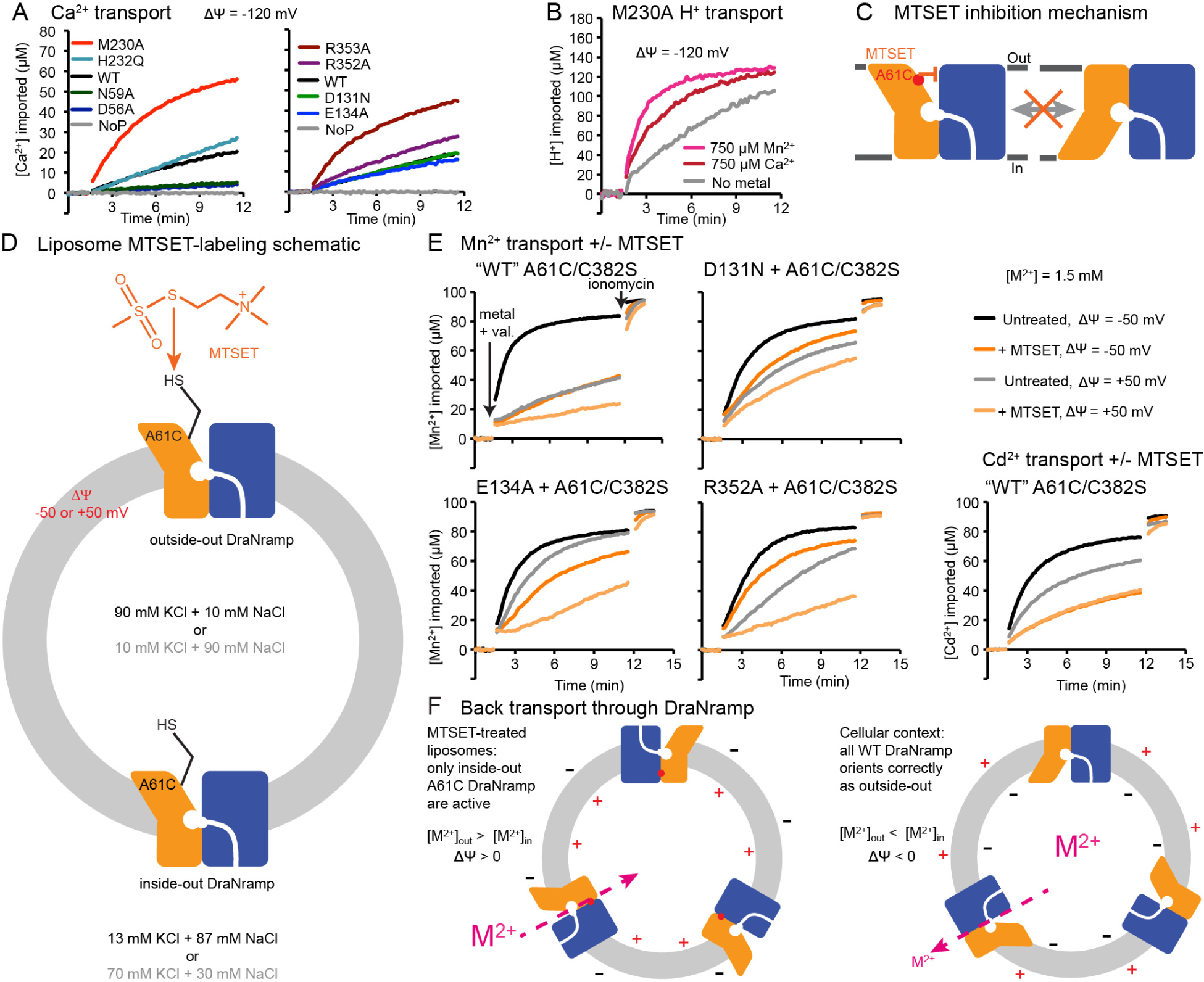
Test of DraNramp back transport. (A) Representative time traces (n ≥ 4) of Ca^2+^ uptake into proteoliposomes. M230A and R353A increased Ca^2+^ transport, D56A and N59A eliminated Ca^2+^ uptake, and other mutants behaved comparably to WT. (B) Representative time traces (n = 4) of Ca^2+^-stimulated H^+^ uptake into proteoliposomes by M230A. (C) Labeling A61C with positively-charged MTSET likely inhibits DraNramp metal transport by locking the protein in an outward-open conformation. (D) Membrane-impermeable MTSET only modifies A61C of DraNramp molecules oriented outside-out after reconstitution into liposomes. Inside-out DraNramp, which we assume is ∼50% of the inserted protein, is protected from modification and thus capable of conformational cycling. (E) Representative metal uptake traces (n = 4) of A61C constructs at ΔΨ = −50 or +50 mV in the presence or absence of MTSET. Interestingly, WT-like A61C/C382S does transport some Mn^2+^ at ΔΨ ≥ 0, unlike the true WT. Mutations to D131, E134, and R352 all increased Mn^2+^-transport at +50 mV even after MTSET labeling. Cd^2+^ transport by WT-like was less dependent on voltage, and showed more residual transport after MTSET labeling. Addition of ionomycin shuttles divalent cations to show the maximum signal. (F) A61C liposome assay mimics cellular context for DraNramp back-transport. Metal influx into MTSET-treated proteoliposomes occurs down a concentration gradient but against a ΔΨ > 0, just as metal efflux would *in vivo*.

**Figure 8—figure supplement 1.**
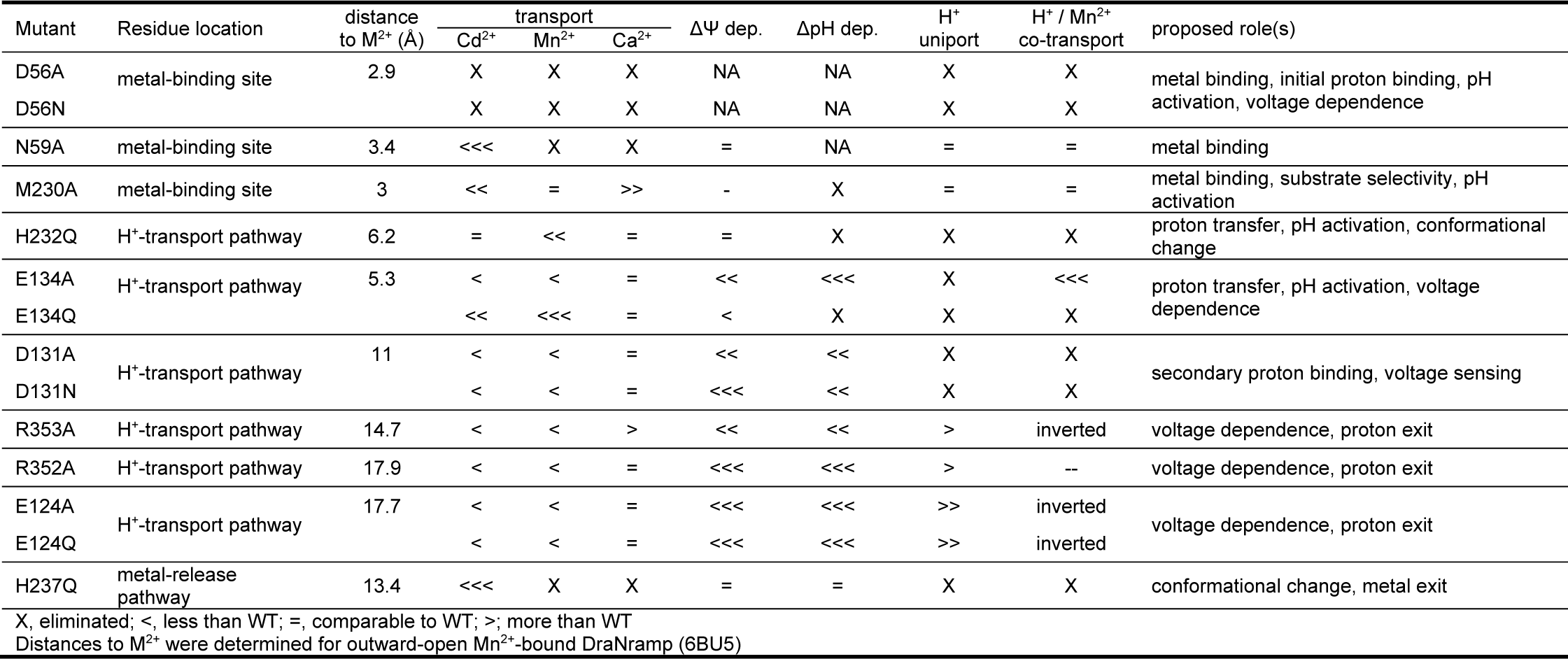
Summary of the results presented in Figures 4-6.

